# Enhanced canonical Wnt signaling during early zebrafish development perturbs the interaction of cardiac mesoderm and pharyngeal endoderm and causes thyroid specification defects

**DOI:** 10.1101/2019.12.19.880815

**Authors:** Isabelle Vandernoot, Benoît Haerlingen, Achim Trubiroha, Pierre Gillotay, Véronique Janssens, Robert Opitz, Sabine Costagliola

## Abstract

**Background:** Congenital hypothyroidism (CH) due to thyroid dysgenesis is a frequent congenital endocrine disorder for which the molecular mechanisms remain unresolved in the far majority of cases. This situation reflects in part our still limited knowledge about the mechanisms involved in the early steps of thyroid specification from the endoderm, in particular the extrinsic signaling cues that regulate foregut endoderm patterning. In this study, we used small molecules and genetic zebrafish models to characterize the role of various signaling pathways in thyroid specification.

**Methods:** We treated zebrafish embryos during different developmental periods with small molecule compounds known to modulate the activity of Wnt signaling pathway and observed effects in thyroid, endoderm and cardiovascular development using whole mount *in situ* hybridization and transgenic fluorescent reporter models. We used an antisense morpholino to create a zebrafish acardiac model. For thyroid rescue experiments, BMP pathway induction in zebrafish embryos was obtained by using heatshock inducible transgenic lines.

**Results:** Interestingly, combined analyses of thyroid and cardiovascular development revealed that overactivation of Wnt signaling during early development leads to impaired thyroid specification concurrent with severe defects in the cardiac specification. When using a model of morpholino-induced blockage of cardiomyocyte differentiation, a similar correlation was observed, suggesting that defective signaling between cardiac mesoderm and endodermal thyroid precursors contributes to thyroid specification impairment. Rescue experiments through transient overactivation of BMP signaling could partially restore thyroid specification in models with defective cardiac development.

**Conclusion:** Collectively, our results indicate that BMP signaling is critically required for thyroid cell specification and identify cardiac mesoderm as a likely source of BMP signals.

## Introduction

The thyroid is an endoderm-derived gland developing from the most anterior part of the gut tube. The thyroid organogenesis starts with the specification of its anlage, a group of thyroid precursor cells that are characterized by co-expression of a unique combination of transcription factors comprising NKX2-1, PAX8, FOXE1, and HHEX (1).

The median anlage develops into a diverticulum that evaginates and loses contact with the ventral foregut endoderm to relocate deeper into the subpharyngeal mesenchyme. Terminal differentiation, which leads to functional follicle formation, is initiated during the migration of the thyroid primordium (2).

Congenital hypothyroidism (CH) is the most frequent congenital endocrine disorder, affecting approximately one of 2500 in human newborns (3). CH is characterized by reduced serum thyroid hormone levels at birth. It is caused in 85% of the cases by thyroid dysgenesis (TD due to ectopy, athyreosis, or hypoplasia, resulting from an aberration of the thyroid gland embryological development. The molecular mechanisms leading to TD are mostly unknown, with mutations in the known thyroid transcription factors explain only 5% of TD cases (1). It probably reflects our limited knowledge about intrinsic factors and external signals coordinating thyroid organogenesis and suggests that unknown genetic and/or epigenetic factors might be crucial for thyroid development (4).

Zebrafish is a valuable model system that has already helped us to improve further our understanding of morphogenetic processes and gene networks involved in thyroid organogenesis (5). Over the past two decades, zebrafish has gained much attention as a genetically tractable vertebrate model to study organogenesis (6). Embryos’ optical clarity allows direct visualization of developmental processes and their pathological deviations. Small size, high fecundity, external fertilization, rapid development, and short generation time are important attributes underscoring the utility of zebrafish study of early developmental processes. The value of zebrafish for studies on thyroid development is supported by the fact that several aspects of thyroid morphogenesis are well conserved between zebrafish and mammals (7), (8). Early morphogenetic events, such as thyroid specification, budding, and relocalization into the subpharyngeal mesenchyme, show many similarities in fish and mouse (reviewed in ref. (9)). Moreover, the developing thyroid expresses a similar, but not identical, set of transcription factors in mouse (*NKX2-1, PAX8, HHEX*, *FOXE1* (1)) and zebrafish (*nkx2.4b* (10), *pax2a*, *pax8*, *foxe1*, *hhex* (7), (10)) embryos. Invalidation of the known thyroid transcription factors in mice leads to athyreosis (*NKX2-1, PAX8, FOXE1, HHEX*) or thyroid ectopy (*FOXE1*) (11). Zebrafish with loss-of-function of *nkx2.4b*, *pax2a,* and *hhex* similarly display athyreosis (7), (9), (12). However, differences exist in the timing of specific morphogenetic events and the anatomy of the mature thyroid tissue. Indeed, although thyroid follicles are encapsulated in a compact organ in mouse, thyroid follicles are loosely scattered along the pharyngeal midline in zebrafish.

In this study, we mainly focused on molecular mechanisms that govern the first steps of thyroid specification. At present, the signals that trigger the specification of pharyngeal endodermal cells into thyroid precursors remains unknown. Recent studies demonstrate that tissue-tissue interactions, in particular with the cardiac mesoderm and pharyngeal blood vessels, are essential for correct thyroid development (13), (5). Defective pharyngeal vessel development has been shown to cause thyroid anomalies in mice and zebrafish (14), (15). There are also pieces of evidence coming from human disease studies since an increased prevalence of congenital cardiovascular anomalies is observed in patients with thyroid dysgenesis compared to the healthy population (16). However, the mechanisms underlying tissue-tissue interactions during thyroid development (particularly between the developing thyroid and the cardiac mesoderm and pharyngeal vessels) and the signals involved in this crosstalk are poorly understood. While a crucial role of FGF expression in the mesoderm surrounding the developing thyroid has been demonstrated in mouse and zebrafish (17), (18), the potential functions of other major signaling pathways in thyroid organogenesis is still to be clarified (2), (19). A significant advantage of the zebrafish model in cardiovascular development studies is that zebrafish embryos and larvae can survive for several days without a functional heart or in the absence of blood circulation (20). This facilitates the characterization of developmental effects over an extended developmental period, compared to most mouse models with cardiovascular defects.

To improve our understanding of the role of extrinsic signaling cues in thyroid development, we recently exploited the amenability of zebrafish embryos for small molecule screenings to identify candidate signaling pathways (12), (21). From a literature review, we first identified a collection of small molecules known to interfere with common signaling pathways in zebrafish. The pharmacological screening readily identified modulators of Wnt, BMP, FGF, and TGF-β signaling to cause distinct effect patterns of disturbed thyroid organogenesis (e.g., agenesis, hypoplasia, ectopy) (21). In this paper, we focused on the phenotypic description, using thyroid and cardiovascular markers, of early thyroid development after Wnt modulation and propose a mechanistic explanation.

Wnt signaling pathway is an evolutionarily conserved signal transduction pathway that regulates crucial aspects of cell fate determination, cell migration, cell polarity and cell differentiation during embryonic development (22). Wnt proteins are secreted glycoproteins that bind to the N-terminal extra-cellular cysteine-rich domain of the Frizzled (Fz) receptor family, interacting with the LRP5/6 co-receptors. To date, several intracellular signaling branches/cascades downstream of the Fz receptors have been identified including a canonical (β-catenin dependent) pathway and a non-canonical (β-catenin-independent) pathway. Some Wnt ligands act preferentially on one pathway or the other, but the activated pathway mostly depends on the cellular context. Without Wnt canonical signaling activation, cytoplasmic β-catenin is degraded by a β-catenin destruction complex, which includes Axin, adenomatosis polyposis coli (APC), protein phosphatase 2A, glycogen synthase kinase 3 (GSK3) and casein kinase 1α (CK1). Binding of a Wnt ligand to its receptor leads to a series of events that disrupt the β-catenin destruction complex, which is required for the β-catenin translocation into the nucleus and activation of target genes (23), (24).

The functions of Wnt/β-catenin signaling in embryogenesis have been extensively studied in different animal models. It plays a crucial biphasic role in heart development, as demonstrated in *in vitro* cells and zebrafish: in pregastrula stages, canonical Wnt promotes the specification of the precardiac mesoderm into cardiomyocyte progenitors, but it inhibits the cardiac differentiation of these cells during gastrula stages (25), (26). It has already been demonstrated that this pathway also plays a very important role in the specification and development of endoderm-derived organs, like liver and pancreas. An anterior-posterior gradient of Wnt activity in *Xenopus* plays a crucial role in endodermal patterning, the anterior endoderm giving rise among others to liver, lung, and thyroid, the posterior endoderm leading to intestinal fate (27). At later stages in zebrafish, Poulain *et al.* have shown that Wnt pathway is necessary for liver progenitors proliferation (28). However, very little data exist on the role of Wnt in thyroid development and/or maintenance.

Therefore, this study aims to examine the effects of the canonical Wnt signaling pathway on thyroid development in zebrafish, in regard to the adjacent cardiogenesis. We used different chemicals and genetic models to modulate canonical Wnt signaling during gastrula and early somitogenesis stages and observed effects of such modulation on thyroid organogenesis. We succeeded to partially rescue thyroid defects obtained after Wnt activation when combining this chemical treatment with induction of BMP pathway using heatshock inducible transgenic lines.

## Material and methods

### Zebrafish husbandry and embryo culture

Zebrafish (*Danio rerio*) embryos were obtained from natural spawning of adult fish, raised at 28.5°C according to Westerfield (29) and staged in hours postfertilization (hpf) as described by Kimmel *et al*.. (30) In addition to wild-type (31) and *casper* mutant lines (32), different transgenic lines were used in this study: *Tg(tg:mCherry*) (5), *Tg(kdrl:EGFP*) (33), *Tg(myl7:EGFP*) (34), *Tg(7xTCF-Xla.Siam:GFP)ia4* (35), *Tg(sox17:EGFP)* (36)*, Tg(hsp70l:bmp2b)* (37) and *Tg(hsp70l:wnt8a-EGFP*) (38). Embryos were dechorionated at 24 hpf using 0.6 mg/mL pronase (Sigma), anesthetized in 0.016% tricaïne (Sigma), fixed in 4% phosphate-buffered paraformaldehyde (PFA; Sigma) overnight at 4°C, washed in PBS containing 0.1% Tween 20 (PBST), gradually transferred to 100% methanol, and stored at −20°C until used for *in situ* hybridization or immunofluorescence analyses. If indicated, pigmentation of embryos was inhibited by adding 0.003% 1-phenyl-2-thiourea (PTU; Sigma) (39) to the embryo medium at 24 hpf. All zebrafish work at the Institute of Interdisciplinary Research in Molecular Human Biology followed protocols approved by the Institutional Animal Care and Use Committee.

### Small molecule treatment

We used timed embryo treatment with BIO (Sigma, B1686), 1-Azakenpaullone [AZA] (Sigma, A3734), and IWR-1 (Sigma, I0161) to modulate the canonical Wnt signaling pathway during distinct developmental periods. BIO and AZA act as canonical Wnt activators due to GSK3β inhibition (40). In contrast, IWR-1 stabilizes the destruction complex, thereby acting as an inhibitor of Wnt signaling (41). Stock solutions of BIO (5 mM), AZA (5 mM), and IWR-1 (10 mM) were prepared in dimethyl sulfoxide (DMSO). Test solutions were prepared by diluting the stock solutions in embryo medium. A control treatment containing 0,1% DMSO was used in all experiments involving a drug treatment.

### Heat-shock treatments

Timed global overactivation of Wnt and BMP signaling was induced by heat-shock treatment of the progeny from matings of WT fish with heterozygous *Tg(hsp70l:wnt8a-GFP*) (38) or *Tg(hsp70l:bmp2b)* (37) fish, respectively. Embryos obtained from these matings were raised under standard conditions until heat-shock to induce global overexpression of EGFP-tagged Wnt8a or Bmp2b. For the heat-shock treatments, embryos were transferred to dishes containing prewarmed embryo medium at 40°C and incubated in an incubator for 30 min at 40°C. After heat-shock treatment, embryos were transferred to Petri dishes containing fresh medium and allowed to develop further at 28.5°C under standard conditions. Carriers of the *hsp70l:wnt8a-EGFP* transgene were identified 3 hours after heat-shock by means of their EGFP expression. Embryos carrying the *hsp70l:bmp2b* transgene were identified by their dorsalized phenotype if heat-shocked at early somitogenesis or by PCR genotyping if embryos were heat-shocked at later stages (15 or 20 hpf). For the latter, PCR genotyping was performed after completion of WISH experiments as previously described (42) using the following primers: Forward 5’-CATGTGGACTGCCTATGTTCATC-3’ (primer located in *hsp70l* promoter sequence); Reverse 5’-GAGAGCGCGGACCACGGCGAC-3’ (primer located in *bmp2b* coding sequence).

### Whole-mount in situ hybridization (WISH)

DNA templates for synthesis of *nkx2.4b, tg, nkx2.5, egfp, mef2cb, gata4, gata5, hhex, foxa2, foxa3, pdx1*, *prox1,* and *bmp4* riboprobes were generated by PCR (see **Supplemental Table 1** for primer sequences). Plasmids for *amhc, vmhc*, *myl7, scl,* and *hand2* riboprobes have been used as described (43), (44), (45). Single-color WISH using digoxigenin (DIG)-labeled riboprobes and anti-DIG antibody conjugated to alkaline phosphatase was performed essentially as described in Thisse and Thisse (46) and Opitz *et al* (5). Riboprobe hybridization was performed at 65°C overnight, and probes were detected using an anti-DIG antibody (1:6000; Roche). Staining reactions were performed with the alkaline phosphatase substrates BM Purple (Roche) or NBT/BCIP (Roche). For dual-color WISH, riboprobes labeled with DIG, dinitrophenol (DNP), or fluorescein (FLU) were used, and sequential alkaline phosphatase staining was performed with BM Purple and Fast Red (Sigma) as described (21).We first detect the DIG or DNP riboprobes with anti-DIG or anti-DNP antibody using BM Purple (for most genes) or NBT/BCIP (for *tg*) staining solution and used an anti-FLU antibody in combination with FastRed for detection of the FLU probe in a second step. Removal of the antibodies after the first staining step was performed by 2x 5 min incubation in 100 mM glycine-HCl (pH 2.2).

Fluorescent WISH (FISH) using a DIG-labeled riboprobe for *nkx2.4b* was performed as described (5). Antibodies used in WISH and FISH experiments are listed in **Supplemental Table 2**. Stained embryos were washed in PBST, postfixed in 4% PFA and embedded in 90% glycerol for whole-mount imaging or in 7% low melting point agarose (Lonza) for vibratome sectioning. Whole-mount images of WISH and FISH were acquired using a MZ16F Leica stereomicroscope equipped with a DFC420C camera or a Leica microscope DMI6000B equipped with a DFC365FX camera, respectively. Vibratome tissue sections at 50–60μm thickness were cut on a Leica VT1000S vibratome and mounted in Glycergel (Dako). Confocal images of vibratome sections were acquired using an LSM510 confocal microscope (Zeiss).

### Whole-mount immunofluorescence (WIF)

WIF was performed essentially as described (47). Briefly, after rehydration into PBST, embryos were immersed in blocking buffer (PBST containing 1.0% DMSO, 1% bovine serum albumin [BSA], 5% horse serum, and 0.8% Triton X-100) for 2 h. Embryos were then incubated overnight in blocking buffer containing primary antibodies at 4°C. After several washing steps in PBST containing 1% BSA, embryos were incubated with secondary antibodies overnight at 4°C. Specifications and sources of primary and secondary antibodies used to detect EFGP green fluorescent protein (48) and pSMAD1/5/8 protein in zebrafish embryos are provided in **Supplemental Table 2**. Stained embryos were washed in PBST and embedded in 90% glycerol for fluorescence microscopy. Images were acquired with a Leica DFC365FX camera mounted on a Leica DMI6000B. Combined FISH and WIF staining was performed as described (5). Confocal images were acquired using a LSM510 confocal microscope (Zeiss).

### RNA extraction and RT-qPCR

For total RNA preparation, pools of 10-20 embryos per sample were lysed in RNeasy Lysis buffer (Qiagen) containing 1% 2-mercaptoethanol (Sigma). Total RNA was isolated using RNeasy RNA preparation microkit (Qiagen) according to the manufacturer’s instructions, including on-column treatment with DNase I (Qiagen). Reverse transcription was done using Superscript II kit (Invitrogen). Reverse transcription quantitative PCR (RT-qPCR) was performed in duplicate using Kapa SYBR Fast (KapaBiosystems) mix and a CFX Connect Real-Time PCR System (Biorad).

Relative values for target transcript abundance in individual samples were determined by the comparative C_T_ method (ΔΔC_T_) according to Pfaffl *et al.* (2001) (49), and data are presented as relative expression values normalized to the reference gene rpl13 (NM_212784.1) whose expression remained constant upon treatment. Primers used were as follows: *rpl13*, forward TCTGGAGGACTGTAAGAGGTATGC, reverse AGACGCACAATCTTGAGAGCAG (50); *egfp*, forward AGAACGGCATCAAGGTGAAC, reverse TGCTCAGGTAGTGGTTGTCG. Gene expression profile was confirmed in triplicate (3 different batches of embryos). Data were expressed as mean ± SD. Pairwise comparisons were performed using the Student *t*-test.

### Fluorescence-activated cell sorting

Cell suspensions were prepared from *Tg(sox17:EGFP)* transgenic embryos at 30 hpf. Embryos were digested in HBSS (Gibco) containing 0.25% trypsin (Gibco) and 2 mM EDTA (Invitrogen). Single-cell solutions were obtained by constant pipetting. Reaction was stopped by adding CaCl2 (final concentration: 1 mM) and FBS (final concentration: 10%). Cells were pelleted by centrifugation and washed with PBS containing 0.4% BSA and 2 mM EDTA. Finally, resuspended cells were passed through a 40 µM nylon mesh (Falcon) into a FACS tube (Falcon) and the GFP+ fractions were analyzed by a fluorescent-activated cell sorter (FACS Aria: FACSDiva Software (BD)). About 50 embryos were sampled per condition, and each experiment was performed in triplicate.

### Morpholino microinjection

For inhibition of the *mef2ca* and *mef2cb* transcripts, zebrafish embryos were injected with morpholino (MO) antisense oligonucleotides that have previously been validated for their knockdown specificity and efficacy (51), (52). 3nL (2ng/nL) of a translation-blocking *mef2d/c*-MO (tb-MO; 5′-ATGGGGAGGAAAAAGATCCAGATTC-3′) was injected as previously described (51). Working solutions of MOs were prepared in 0.12 M KCl containing phenol red, and 3 nL were microinjected into the yolk of one- to two-cell stage embryos. Non-injected embryos served as control embryos.

### Statistics

For the statistical analyses of thyroid and cardiac phenotypes in the rescue experiments, Fisher tests were conducted using the software package GraphPad Prism 4.0 (GraphPad, San Diego, CA). Differences were considered significant at p < 0,05.

## Results

### Drug-induced manipulation of canonical Wnt signaling disrupts early thyroid development

We recently performed a small molecule screen to identify signaling pathways involved in early zebrafish thyroid development (21). In these experiments, pharmacological manipulations of canonical Wnt signaling during gastrula stage resulted in abnormal thyroid marker expression, and most notably, in a severe reduction of *nkx2.4b* expression at thyroid anlage stages (28 hpf) following overactivation of Wnt signaling.

To characterize in more detail the thyroidal effects resulting from enhanced Wnt signaling, we performed additional treatment experiments with BIO and AZA, two drugs known to activate Wnt/β-catenin signaling (40). Consistent with our previous results, treatment of zebrafish embryos with BIO or AZA during the gastrulation period (6 to 10 hpf) caused concentration-dependent decreases of *nkx2.4b* expression in 28 hpf embryos (**Figure 1**). Maximal effects on thyroidal *nkx2.4b* expression (i.e. complete absence of a detectable WISH staining) were evident at 5 µM BIO and 5 µM AZA. In addition to the marked effects on the thyroidal *nkx2.4b* expression domain, BIO and AZA treatments also caused concentration-dependent decreases in the size of the *nkx2.4b* expression domain in the forebrain. The drug-induced effects on forebrain *nkx2.4b* expression correlated closely with other visible signs of global posteriorization, including loss of anterior neural tissue and reduced size or absence of the eyes (53), (54). Although we did not quantify the relative size reductions of thyroidal and forebrain *nkx2.4b* expression domains in 28 hpf embryos following different BIO and AZA treatments, we noted a similar sensitivity to drug treatment for both expression domains when embryos were treated between 6 – 10 hpf.

**Fig. 1.**
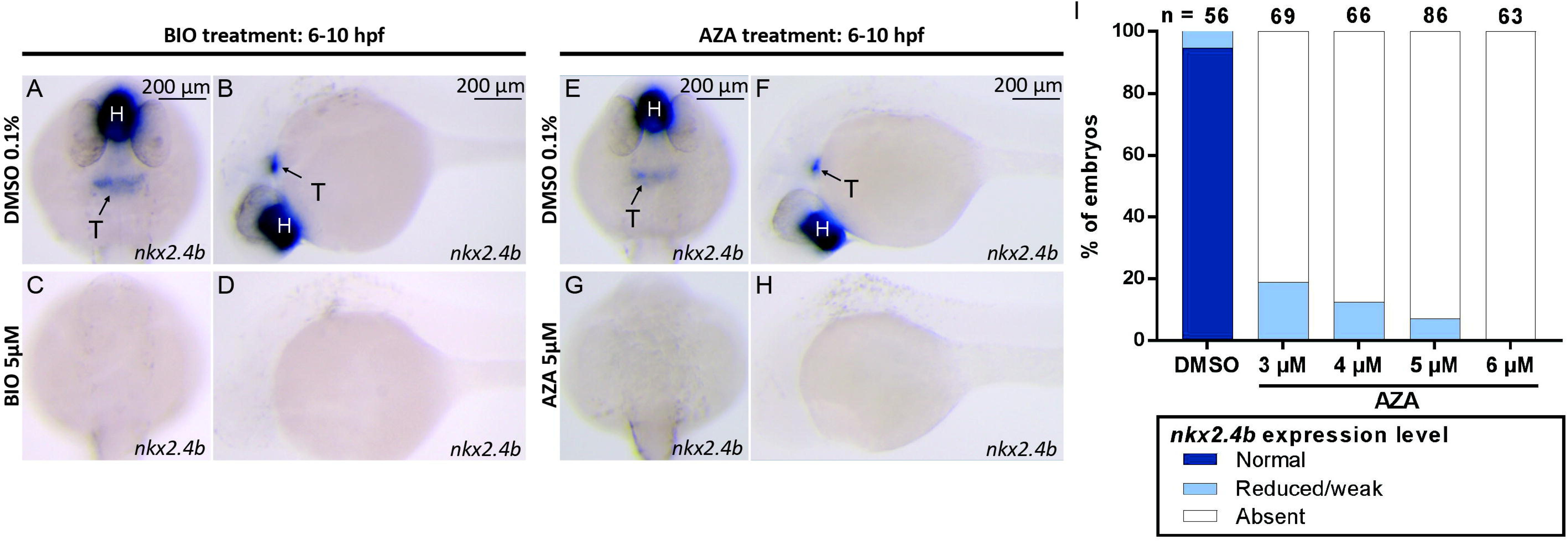
Drug-induced overactivation of canonical Wnt signaling during gastrulation impairs thyroid specification. (**A-H**) Expression of the early thyroid marker *nkx2.4b* in 28 hpf embryos following treatment with 0.1% DMSO (vehicle control) and two Wnt-inducing drugs, BIO and azakenpaullone (AZA) between 6 and 10 hpf. In control embryos (A,B,E,F), *nkx2.4b* is expressed in the thyroid anlage (T) and the ventral forebrain (hypothalamus, H). In response to BIO (C-D) and AZA (G-H) treatment, *nkx2.4b* mRNA expression is lost in both domains in the majority of embryos. Arrows point to *nkx2.4b* expression in the thyroid anlage. Dorsal views with anterior oriented to the top (A,C,E,G) and lateral views with anterior oriented to the left (B,D,F,H) are shown. (**I**): Quantification of the proportion of specimen displaying thyroid specification defects following AZA treatment, as determined by *nkx2.4b* staining of 28 hpf embryos. Results are presented as the percentage of embryos displaying a particular phenotype, showing a concentration-dependent *nkx2.4b* mRNA loss of expression. The total number of specimens analyzed for each treatment group is provided on the top of each column. Scale bar, 200 µm.

Concentration-dependent effects on thyroid development were also evident when BIO- and AZA-treated embryos were analyzed for thyroid marker expression at 55 hpf (**Figure 2**). WISH analysis of *nkx2.4b* expression revealed concurrent reductions in the size of thyroidal and forebrain expression domains in treated embryos. Moreover, BIO and AZA treatment strongly reduced the expression of the functional thyroid differentiation marker, thyroglobulin (*tg),* in 55 hpf embryos. However, we also noted that the majority of 55 hpf embryos treated with high drug concentrations during gastrula stages displayed at least some residual staining for *nkx2.4b* and *tg* despite undetectable thyroid marker expression at the thyroid anlage stage (28 hpf). This was observed in all experiments involving drug treatment from 6 to 10 hpf.

**Fig. 2.**
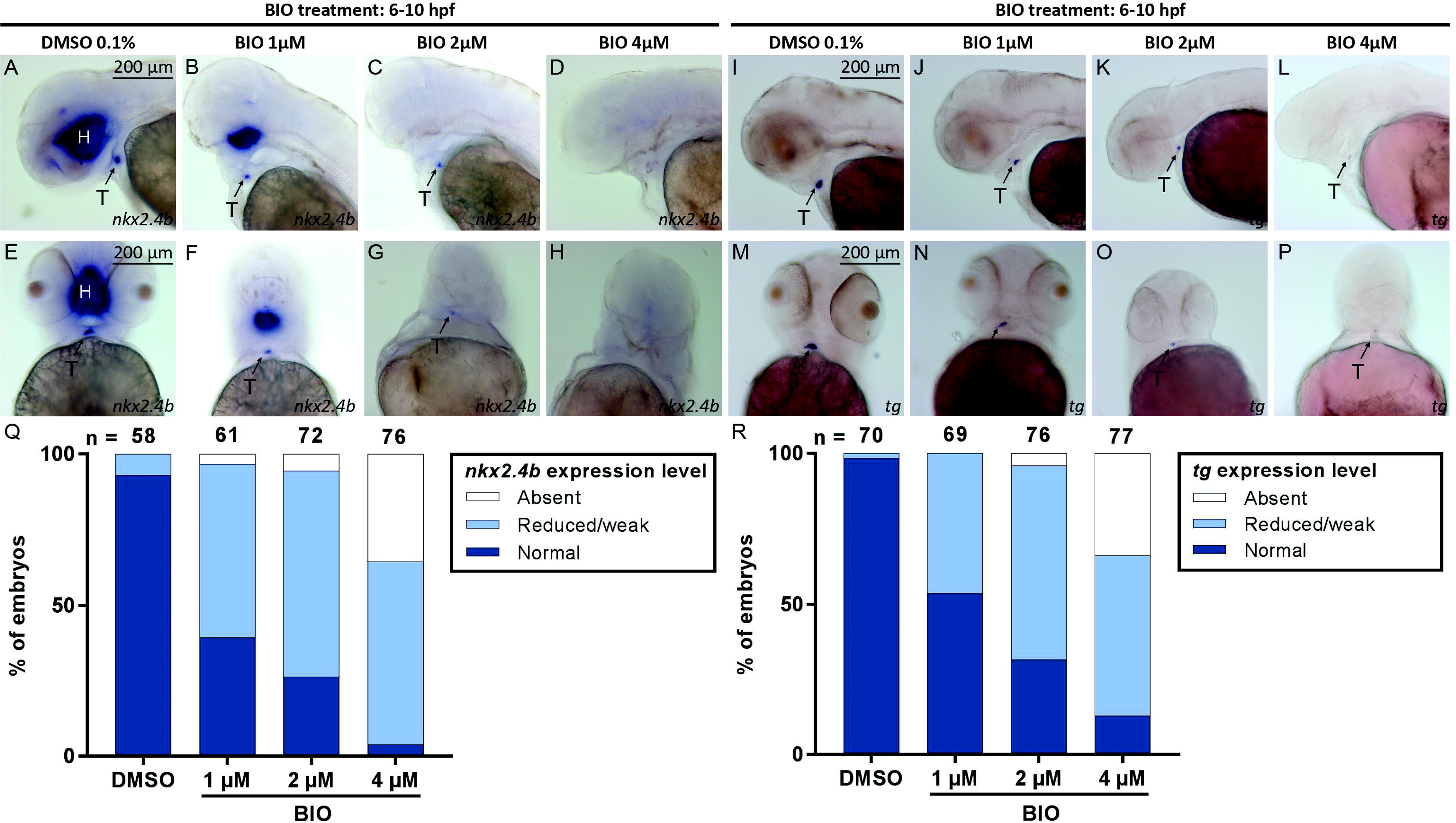
Drug-induced overactivation of canonical Wnt signaling during gastrulation impairs the thyroid primordium formation. (**A-H**) Expression of the thyroid marker *nkx2.4b* in 55 hpf embryos following treatment with 0.1% DMSO (vehicle control) and the Wnt-inducing drug BIO between 6 and 10 hpf. In control embryos (A,E), *nkx2.4b* is expressed in the thyroid primordium (T) and the ventral forebrain (hypothalamus, H). In response to BIO (B-D,F-H) treatment, *nkx2.4b* mRNA expression is reduced or lost in both tissues in a concentration-dependent manner. Arrows point to *nkx2.4b* expression in the thyroid primordium. Lateral views with anterior oriented to the left (A-D) and dorsal views with anterior oriented to the top (E-H) are shown. (**I**): Quantification of the percentage of embryos displaying a particular phenotype following BIO treatment, as determined by *nkx2.4b* staining of 55 hpf embryos. Total numbers of specimens analyzed for each treatment group are provided on the top of each column. (**J-Q**) Expression of the terminal thyroid differentiation marker *tg* in 55 hpf embryos following treatment with 0.1% DMSO (vehicle control) and the Wnt-inducing drug BIO between 6 and 10 hpf. In control embryos (J,N), *tg* is exclusively expressed in the thyroid primordium (T). In response to BIO treatment (K-M,O-Q), *tg* mRNA expression is reduced or lost in a concentration-dependent manner. Arrows point to the thyroid primordium. Lateral views with anterior oriented to the left (J-M) and dorsal views with anterior oriented to the top (N-Q) are shown. (**I**): Quantification of the percentage of embryos displaying a particular phenotype following BIO treatment, as determined by *tg* staining of 55 hpf embryos. Total numbers of specimens analyzed for each treatment group are provided on the top of each column. Scale bar, 200 µm.

Conversely, treatment of gastrulating embryos with 10 µM IWR-1, a small molecule inhibitor of canonical Wnt signaling, resulted in additional ectopic domains of thyroidal *nkx2.4b* and *tg* expression **(Supplementary figure 1).** In 28 hpf embryos, the additional ectopic *nkx2.4b* expression was detected posterior to the orthotopic thyroidal *nkx2.4b* expression domain, whereas in 55 hpf embryos, supernumerary clusters of *tg* expression were detectable at irregular posterior positions. Taken together, these data indicate that the canonical Wnt pathway negatively regulates thyroid specification during gastrulation stage in zebrafish as drug-mediated activation or inhibition of Wnt activity during gastrulation impairs or amplifies thyroid specification, respectively.

### Small molecule compounds rapidly act on Wnt signaling

We next took advantage of an available Wnt signaling biosensor line, *Tg(7xTCF*-*Xla.Siam:GFP)^ia4^*, to verify that the small molecule compounds BIO, AZA and IWR-1 effectively alter canonical Wnt signaling activities within a short space of time under the experimental conditions that resulted in irregular thyroid development. For this purpose, we treated *Tg(7xTCF-Xla.Siam:GFP)^ia4^* embryos with BIO (5 µM), AZA (5 µM) or IWR-1 (10 µM) from 6 to 10 hpf and monitored the expression of *GFP* mRNA during the course of drug treatment by WISH and RT-qPCR **(Supplementary figure 2)**.

WISH analyses of *GFP* mRNA expression in BIO- and AZA-treated embryos showed a rapid and robust up-regulation of *GFP* mRNA expression when compared to DMSO-treated embryos (0.1% DMSO) indicating effective overactivation of canonical Wnt signaling in drugged embryos within 2 hours after the beginning of drug treatment. Consistently, IWR-1 treatment resulted in reduced *GFP* mRNA expression, evident within 2-3 hours after the beginning of drug treatment. Although several embryos treated with these small molecule compounds experienced slight developmental delay, the drug-induced alterations in reporter expression were robustly detectable in comparison with vehicle control embryos. In further experiments, we also corroborated the effects of BIO treatment on *GFP* mRNA expression by RT-qPCR. As shown in **Supplementary figure 2**, treatment of *Tg(7xTCF-Xla.Siam:GFP)^ia4^*embryos with 5 µM BIO resulted in 2.5- to 3-fold increases of whole embryo expression levels of *GFP* mRNA within 2 hours after initiation of BIO treatment. Together, these data demonstrate that, under our experimental conditions, the drug treatments faithfully altered canonical Wnt signaling.

In order to provide an independent line of evidence that defective thyroid development is due to transient overactivation of Wnt/β-catenin signaling during gastrulation, we manipulated Wnt/β-catenin signaling by heat-shock-induced overexpression of *wnt8a* in *Tg(hsp70l:wnt8a-GFP)* embryos (38). *Wnt8a* specifically activates Wnt/β-catenin signaling in zebrafish embryos and a 30 min heat-shock of transgenic *Tg(hsp70l:wnt8a-GFP)* embryos at 6 hpf resulted in a marked neural posteriorization phenotype and disturbed early thyroid development (**Figure 3**). Non-transgenic embryos showed no discernible phenotypes in response to the heat-shock treatment. The thyroid phenotype caused by transient *wnt8a* overexpression in the genetic model closely resembled the phenotypes observed following treatment with 4-5 µM BIO or AZA. Specifically, our WISH analyses failed to detect *nkx2.4b* expression in the thyroid anlage region of heat-shocked embryos carrying the *wnt8a-GFP* transgene at 28 hpf and a detectable, though strongly diminished, *tg* expression in 55 hpf transgenic embryos (**Figure 3**). Taken together, these experiments verified that the selected drug concentrations cause overactivation of Wnt/β-catenin signaling and that enhanced Wnt/β-catenin signaling during gastrula stages leads to defects in thyroid development.

**Fig. 3.**
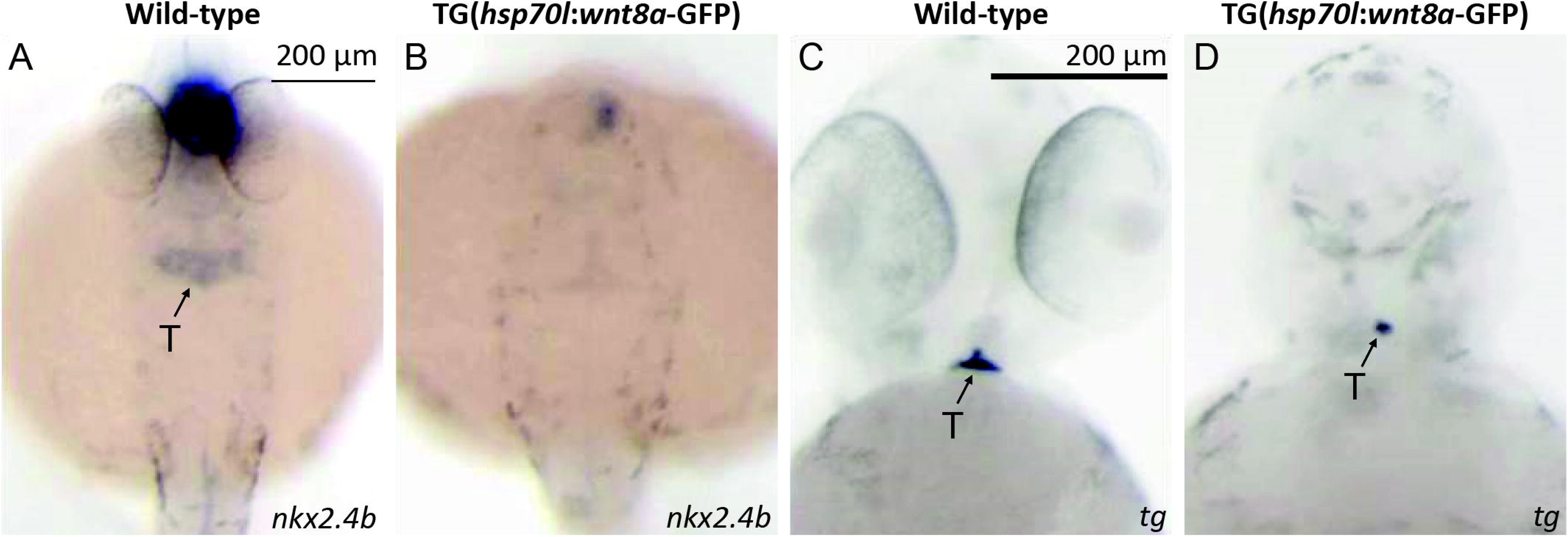
Heat shock-induced overexpression of *wnt8a-GFP* during gastrulation impairs thyroid specification and thyroid primordium formation. (**A,B**) Expression of the thyroid marker *nkx2.4b* in 28 hpf embryos following heat shock (HS) treatment of *Tg(hsp70l:wnt8a-GFP)* embryos at 6 hpf. In response to HS, embryos carrying the HS-inducible *wnt8a-GFP* transgene showed a great reduction or even loss of *nkx2.4b* mRNA expression in the thyroid anlage (T) and the ventral forebrain (hypothalamus, H), whereas expression of *nkx2.4b* in both tissues was unaffected after HS of non-transgenic siblings. Dorsal views with anterior oriented to the top are shown. Arrows point to *nkx2.4b* expression in the thyroid anlage. (**C,D**) Expression of the terminal thyroid differentiation marker *tg* in 55 hpf embryos following heat shock (HS) treatment of *Tg(hsp70l:wnt8a-GFP)* embryos at 6 hpf. Note the strongly reduced size of the thyroid primordium in embryos carrying the HS-inducible *wnt8a-GFP* transgene compared to normal thyroid size in non-transgenic siblings. Similar to drug-induced wnt overactivation, HS-induced overexpression of the *wnt8a-GFP* transgene resulted in a dramatic neural posteriorization phenotype (loss of anterior neural tissue, lack of eyes). Ventral views with anterior oriented to the top are shown. Arrows point to *tg* expression in the thyroid primordium. Scale bar, 200 µm.

### Effects of early Wnt overactivation on endoderm development

Our initial experiments with BIO and AZA showed that treatment with different drug concentrations during gastrulation causes thyroid abnormalities and a strong posteriorization of neural ectoderm. Considering that the latter phenotype is a well-characterized effect of enhanced Wnt signaling during early development (55), we next examined whether endodermal development is similarly affected by a global posteriorization activity due to enhanced Wnt signaling and might thus explain the loss of thyroid marker expression in the anterior endoderm.

To address this question, we first treated embryos of the transgenic *Tg(sox17:EGFP)* line with BIO (5 µM) or AZA (5 µM) between 6 – 10 hpf and analyzed the gross morphology of their foregut endoderm at different embryonic stages, in comparison to that of DMSO-treated embryos . While a neural posteriorization phenotype (loss of anterior neural tissue) was readily detectable in BIO- and AZA-treated *Tg(sox17:EGFP)* embryos, the gross morphology of the anterior endoderm appeared unaffected in these embryos **(Supplementary figure 3)**. In addition, we used FACS to compare the number of GFP+ cells in 30 hpf *Tg(sox17:EGFP)* embryos following treatment with DMSO (0.1%) and BIO (5µM) but did not detect a decrease in the number of GFP+ cells in BIO-treated embryos **(Supplementary figure 3)**. Thus, in contrast to the overt effects on anterior neural tissue development, no visible dysgenesis of the anterior endoderm was detected in *Tg(sox17:EGFP)* following early Wnt overactivation.

To further study possible global endodermal patterning defects in BIO and AZA-treated embryos, we analyzed the expression of a panel of informative endodermal marker genes by WISH (**Figure 4**). We first confirmed that thyroidal expression of *nkx2.4b* and *hhex* is affected similarly by BIO and AZA treatment (see **Figure 4A-D**). In addition to the thyroid, *hhex* is expressed in the developing liver and pancreas. Interestingly, *hhex* expression was selectively repressed in the thyroid anlage of BIO-treated embryos and was maintained in the liver/pancreas region (**Figure 4D**). Notably, the shape and position of the hepatic and pancreatic *hhex* expression domains were altered in BIO-treated embryos. Therefore, we analyzed the expression of *foxa2*, a master regulator of foregut endoderm patterning. Notably, our WISH analyses did not reveal gross changes in the *foxa2* anterior-posterior expression pattern following BIO treatment (**Figure 4E,F**). However, when analyzing *foxa3* expression, a key regulator of posterior endoderm development, we noted that its expression domain appeared extended more anteriorly in many embryos after BIO treatment (**Figure 4G,H**). Analyses of *pdx1* and *prox1a* expression confirmed that hepatic and pancreatic markers are unaffected in BIO-treated embryos and that diminished expression in response to canonical Wnt overactivation is limited to thyroidal marker genes (**Figure 4I-L**).

**Fig. 4.**
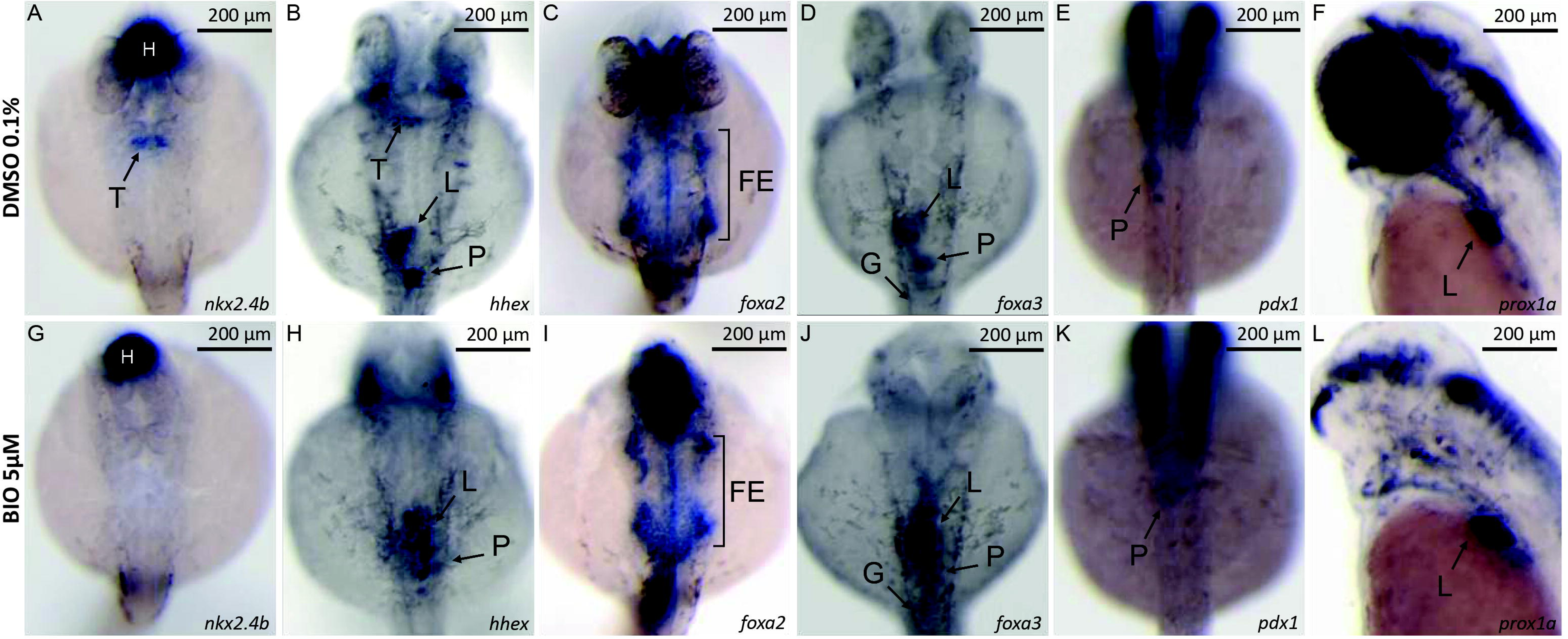
Overactivation of canonical Wnt signaling during gastrula stages has limited effects on markers of endodermal patterning and endodermal organogenesis. (**A-L**) Comparative whole-mount *in situ* hybridization analysis of mRNA expression patterns for a panel of markers of endodermal patterning in embryos treated with 0.1% DMSO (vehicle control) and 5 µM BIO from 6 – 10 hpf. In contrast to the lack of *nkx2.4b* (**A,B**) and *hhex* (**C,D**) expression in the region of the thyroid anlage (T) of BIO-treated embryos, *hhex* expression is maintained in the region of the prospective liver (L) and pancreas (P) development. Robust *foxa2* expression (**E,F**) in BIO-treated embryos indicates that specification of foregut endoderm (FE) was not blocked by Wnt overactivation. However, expression of the mid-/hindgut marker *foxa3* (**G,H**) was expanded towards more anterior regions in several BIO-treated embryos suggesting possible posteriorization effects on endodermal patterning. Importantly, BIO-treated embryos displayed robust staining for all markers of hepatic and pancreatic development including *hhex* (**C,D**), *foxa3* (**G,H**), *pdx1* (**I,J**) and *prox1a* (**K,L**) while showing a specific reduction or lack of markers of early thyroid development. Long arrows point to the thyroid, short arrows to the liver, and arrowheads to pancreatic domains. Brackets in panels **E,F** demarcate the foregut endoderm (FE) domain of *foxa2* expression. Panels A-J show dorsal views with anterior oriented to the top. Panels K and L show lateral views with anterior oriented to the left. Scale bar, 200 µm.

Collectively, our analyses indicate that the loss of thyroid marker expression caused by overactivation of canonical Wnt signaling during gastrulation is not due to a general defect in anterior foregut formation. However, as indicated by the irregular anterior *foxa3* expression in BIO-treated embryos and the posterior extension of thyroid marker expression in IWR-1-treated embryos, we cannot rule out that a Wnt-induced foregut endoderm posteriorisation might contribute to the thyroid defects seen in the most anterior part of the foregut.

### Cardiac phenotype is correlated with thyroid phenotype after gastrula Wnt activation

A recurrent theme in the development of endoderm-derived organs is the critical role of tissue-tissue interactions, particularly between the endoderm and adjacent mesodermal tissues (56), (28). Observations made during our initial small molecule screening experiments (21) indicated that overactive Wnt signaling might induce the concurrent cardiac and thyroidal maldevelopment. To examine this possible relationship in more detail, we first confirmed that drug-induced overactivation of Wnt signaling during gastrula stages results in a severe inhibition of heart formation **(Supplementary figure 4)** as previously reported for genetic models with overactivation of canonical Wnt signaling (26), (25). When treated between 6 to 10 hpf, effects of Wnt-activating drugs on heart development appeared concentration-dependent, and we noticed that development of ventricular cardiomyocytes was more severely affected, compared to that of atrial cardiomyocytes (**Supplementary figure 4**). Notably, the concentrations at which BIO and AZA caused severe cardiac maldevelopment (**Figure 5**) overlapped the range at which repression of thyroid markers was detected.

**Fig. 5.**
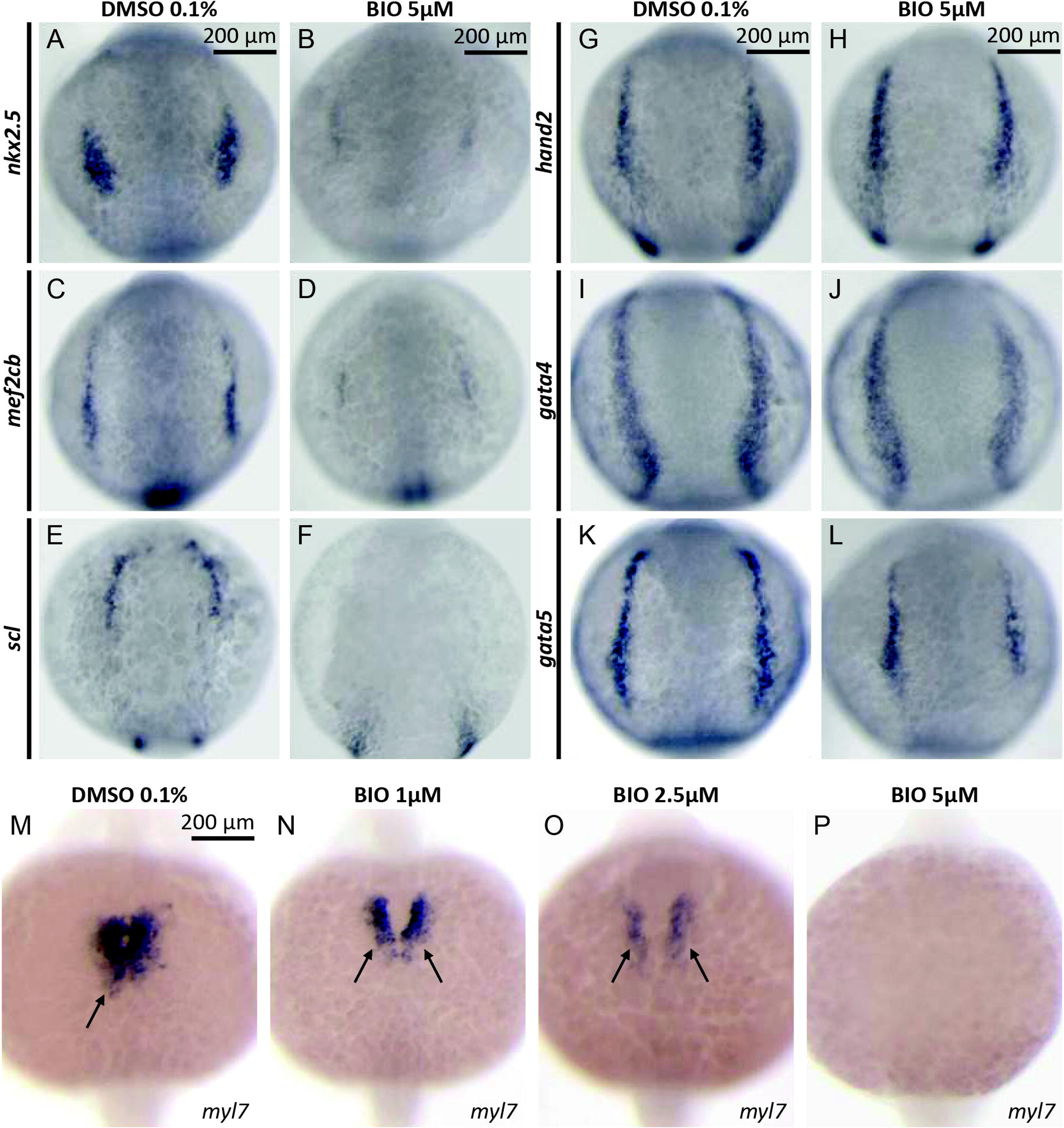
Drug-induced overactivation of canonical Wnt signaling during gastrulation impairs early stages of cardiac development. (**A-L**) Whole-mount *in situ* hybridization analysis of mRNA expression for a panel of cardiac differentiation markers in 13 hpf embryos treated with 0.1% DMSO (vehicle control) and 5 µM BIO from 6 – 10 hpf. BIO treatment severely blocked early stages of cardiac development, as shown by the dramatic reduction in early cardiac markers *nkx2.5* (**A,B**) and *mef2cb* (**C,D**). In addition, there is a severe impairment of endothelial precursor formation, as indicated by the loss of *scl* expression (**E,F**) in BIO-treated embryos. Wnt overactivation specifically affected cardiac and endothelial precursor development, as shown by robust expression of other markers of lateral plate mesoderm development including *hand2* (**G,H**), *gata4* (**I,J**), and *gata5* (**K,L**). Dorsal views with anterior oriented to the top are shown. (**M-P**) Expression analysis of the myocardial marker *myl7* at 19 hpf reveals concentration-dependent effects of early BIO treatment on heart morphogenesis, as evident from the impaired cardiac cone formation, delayed midline fusion of bilateral cardiac fields, and severe depletion of differentiated cardiomyocytes. Arrows point to *myl7* expression in the region of the forming cardiac cone. Dorsal views with anterior oriented to the top are shown. Scale bar, 200 µm.

To characterize the evolution of the cardiac differentiation defects in drugged embryos, we examined the expression of various cardiac differentiation markers at earlier stages of cardiac development (**Figure 5**). Comparative WISH analyses of BIO- and DMSO-treated embryos showed severely reduced expression of transcription factors regulating cardiovascular development such as *nkx2.5*, *mef2cb,* and *scl* in response to Wnt overactivation. Consistent with previous studies employing genetic zebrafish models (25), expression levels of these genes were dramatically decreased as early as 13 hpf in BIO-treated embryos. Conversely, we observed only mild changes, if any, in the expression of transcription factors with a broader lateral plate mesoderm expression domain, including *gata4*, *gata5,* and *hand2* (**Figure 5**). Given that endodermal progenitors of the thyroid lineage develop in close vicinity to the anterior lateral plate mesoderm containing cardiac precursors zebrafish (17), we hypothesized that diminished availability of cardiac mesoderm-borne signaling cues contributes to the defective thyroid development in BIO-treated embryos.

To evaluate a possible causal relationship between inhibited cardiac development and thyroid dysgenesis, we examined the co-occurrence of cardiac and thyroid developmental defects over a range of drug concentrations by dual-color WISH of cardiac (*myl7*) and thyroid (*nkx2.4b*) marker expression in 28 hpf embryos (**Figure 6**). For embryos treated with increasing concentrations of BIO from 6 to 10 hpf, we observed a gradual loss of cardiomyocytes correlated with a gradual reduction in thyroid marker expression (**Figure 6A-F**). Notably, although we observed embryos without any detectable thyroidal *nkx2.4b* expression but still displaying remnant *myl7* positive cardiac tissue, we never observed any embryo with detectable thyroid marker expression in the complete absence of *myl7* cardiac expression. While these observations are compatible with the hypothesis that the availability of cardiac mesoderm and associated signaling cues are prerequisites for thyroid lineage specification to occur, the concurrent global posteriorization phenotype of the embryos prevented us from distinguishing the relative impact of cardiac dysgenesis against a possible posteriorized patterning in the etiology of the thyroid specification defect.

**Fig. 6.**
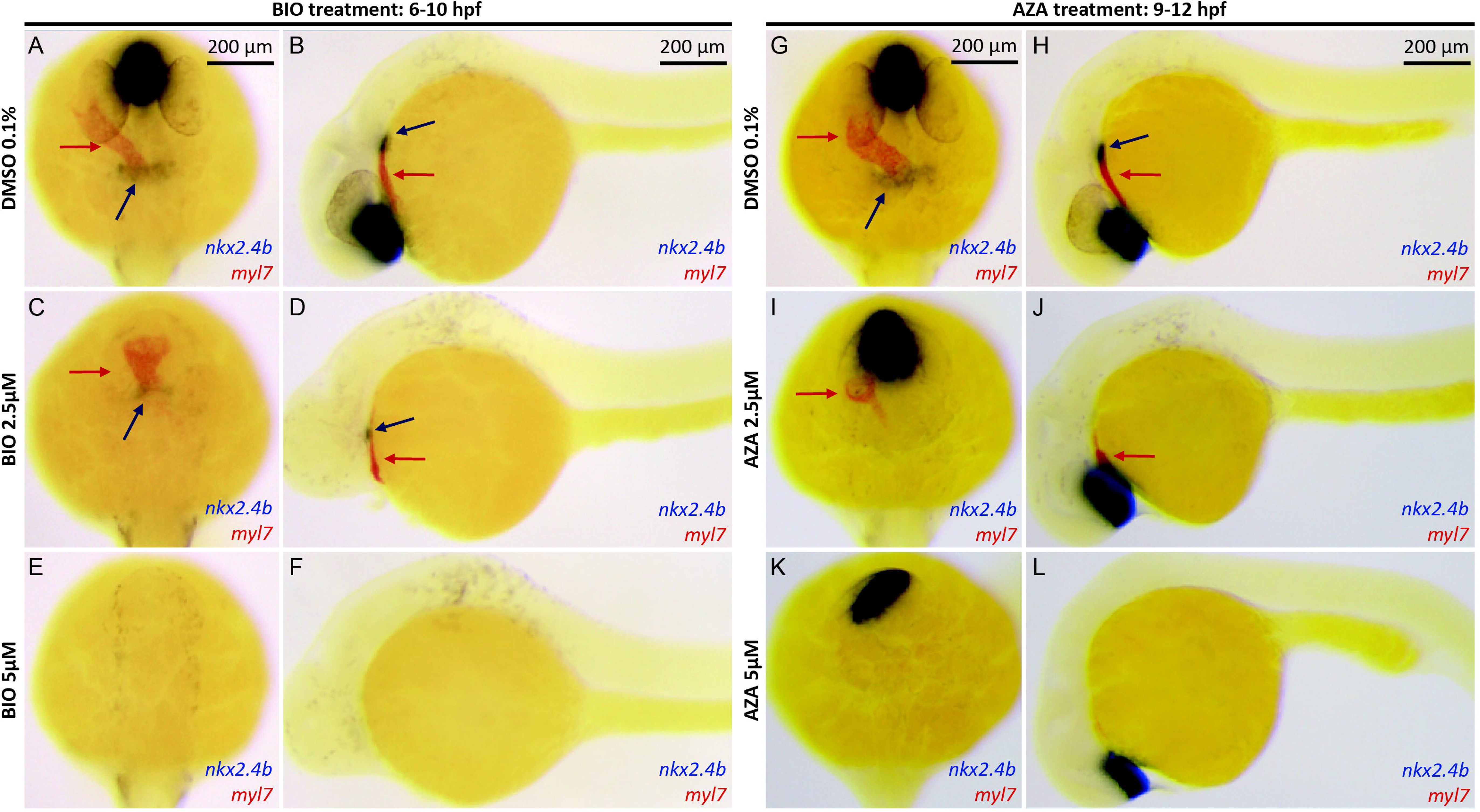
Thyroid specification defects coincide with impaired cardiac development in embryos following drug-induced overactivation of canonical Wnt signaling. (**A-F**) Dual-color whole-mount *in situ* hybridization analysis of mRNA expression of markers for early thyroid (*nkx2.4b*) and cardiac (*myl7*) development in 28 hpf embryos treated with 0.1% DMSO (vehicle control) and varying concentrations of BIO from 6-10 hpf. Wnt overactivation during this period concurrently affected thyroid and cardiac development though thyroid specification appeared slightly more sensitive to BIO treatment. The severity of the thyroid defects was tightly correlated to the degree of neural posteriorization effects. (**G-L**) Dual-color *in situ* hybridization analysis of *nkx2.4b* and *myl7* expression in 28 hpf embryos treated with 0.1% DMSO (vehicle control) and varying concentrations of azakenpaullone (AZA) from 9-12 hpf. Wnt overactivation during this slightly later period resulted in few if any neural posteriorization phenotypes, whereas thyroid and cardiac development were concurrently affected, showing a similar sensitivity to perturbed Wnt signaling. Dorsal views (**A,C,E,G,I,K)** with anterior to the top and lateral views (**B,D,F,H,J,L**) with anterior to the left are shown. Blue arrows point to thyroidal *nkx2.4b* expression and red arrows to *myl7* expression in the heart tube. Scale bar, 200 µm

It has been demonstrated that the later the embryos are treated with Wnt activators during gastrulation, the weaker is the posteriorization effect. Still, cardiac development remains highly sensitive to these later treatments (25). Therefore, we progressively shifted the treatment periods towards later stages and assessed the impact of drug-induced Wnt signaling on global posterization and cardiac and thyroid development in these embryos. As expected, gross morphological signs of posterization were greatly reduced if embryos were treated from 9 to 12 hpf (**Figure 6G-L**). However, Wnt overactivation at these later stages caused still a severe inhibition of cardiac development. Moreover, It appeared that the prevalence and severity of cardiac dysgenesis and thyroid misspecification correlated closely in embryos treated with Wnt-activating drugs between 9 and 12 hpf. We made the same observations when we treated the embryos from 10 to 13 hpf or from 9 to 24 hpf (**Figure 7**). However, we noted that initiation of drug treatment at 11 hpf or later stages caused progressively milder effects on both cardiac and thyroid differentiation as assessed by WISH for cardiac *myl7* expression and thyroidal *nkx2.4b* expression (**Figure 7I,J**).

**Fig. 7.**
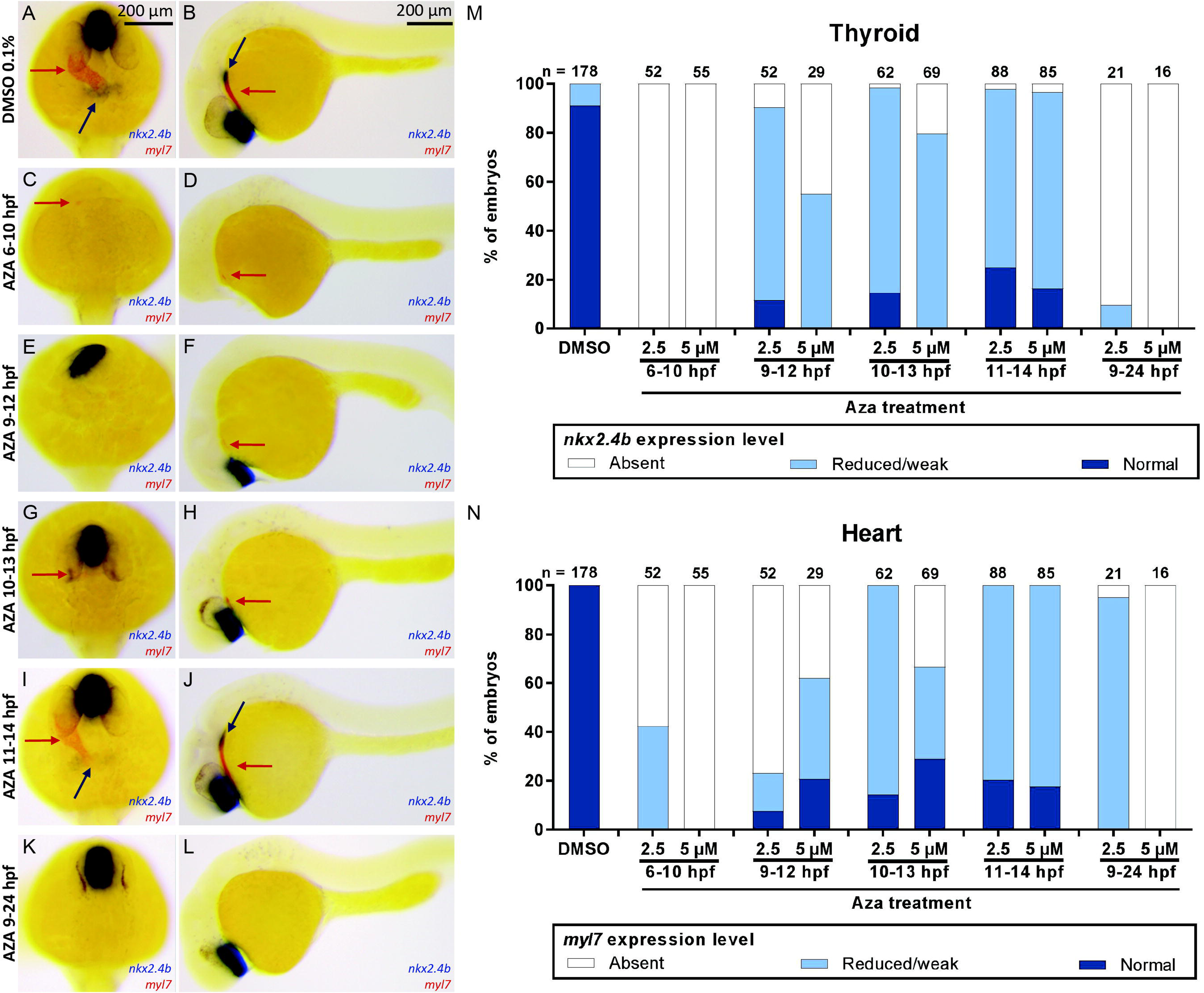
Time-dependent effects of Wnt overactivation on thyroid and cardiac development during gastrulation and early somitogenesis stages. (**A-L**) Dual-color whole-mount *in situ* hybridization analysis of mRNA expression of markers for early thyroid (*nkx2.4b*) and cardiac (*myl7*) development in 28 hpf embryos treated with 0.1% DMSO (vehicle control) and azakenpaullone (AZA) during different periods of early development. Note that neural posteriorization phenotypes became much less prominent and severe when drug-induced Wnt overactivation was initiated at late gastrula stages. Concurrent defects in thyroid and cardiac development were detected for all drug treatments that started before 11 hpf. Prevalence and severity of thyroid and cardiac phenotypes showed a close correlation across different treatment periods. Blue arrows point to thyroidal *nkx2.4b* expression and red arrows to *myl7* expression in the heart tube. Dorsal views (**A,C,E,G,I,K)** with anterior to the top and lateral views (**B,D,F,H,J,L**) with anterior to the left are shown. Scale bar, 200 µm. (**M,N**) Quantification of the proportion of specimen displaying defects in thyroid specification (**M**) and cardiac development (**N**) following AZA treatment, as determined by *nkx2.4b* (**M**) and *myl7* (**N**) staining of 28 hpf embryos. Results are presented as the percentage of embryos displaying a particular phenotype. The total number of specimens analyzed for each treatment group is provided on the top of each column.

### Experimental depletion of cardiomyocyte developmental is associated with thyroid dysgenesis

To further examine the hypothesis that depletion of cardiac mesoderm results in a failure of thyroid cell specification, we sought for cardiac differentiation-defective zebrafish models. To the best of our knowledge, a zebrafish model completely lacking cardiac cells while maintaining normal anterior endoderm has not yet been described. Hinits *et al.* recently described a severe deficiency in cardiomyocyte differentiation in a dual loss-of-function model for zebrafish cardiac transcription factors *mef2ca* and *mef2cb* (51), (52). For our study, we used an antisense morpholino (*mef2c/d*-MO) developed by the same group that reportedly ablates several Mef2 proteins and faithfully replicates the cardiac phenotype present in *mef2ca*/*mef2cb* double mutant fish (52). In our hands, injection of the *mef2c/d*-MO into wild-type embryos, hereafter called Mef2c-deficient embryos, robustly inhibited cardiomyocyte development as evident from only faint *myl7* staining of small and very thin heart tube remnants at 28 and 55 hpf (**Figure 8**). Compared to control embryos, we also observed a greatly reduced expression of *nkx2.4b* in the thyroid anlage of 28 hpf *mef2c/d*-morphant embryos (**Figure 8A-D**). Across several *mef2c/d*-MO injection experiments, we observed that prevalence and severity of cardiac malformation correlated with the extent of impaired thyroid marker expression in 28 hpf morphant embryos. Detectable domains of *nkx2.4b* expression displayed a weaker staining intensity and were of a smaller size in Mef2c-deficient embryos relative to control embryos. Comparison of control and Mef2c-deficient embryos at 55 hpf revealed an irregular organization of *tg*-expressing cells in Mef2c-deficient embryos, which displayed a fainter *tg* staining of individual cells. Collectively, observations made in this Mef2c-deficient zebrafish model of impaired cardiomyocyte differentiation support the contention that thyroid anlage specification relies on normal cardiac development.

**Fig. 8.**
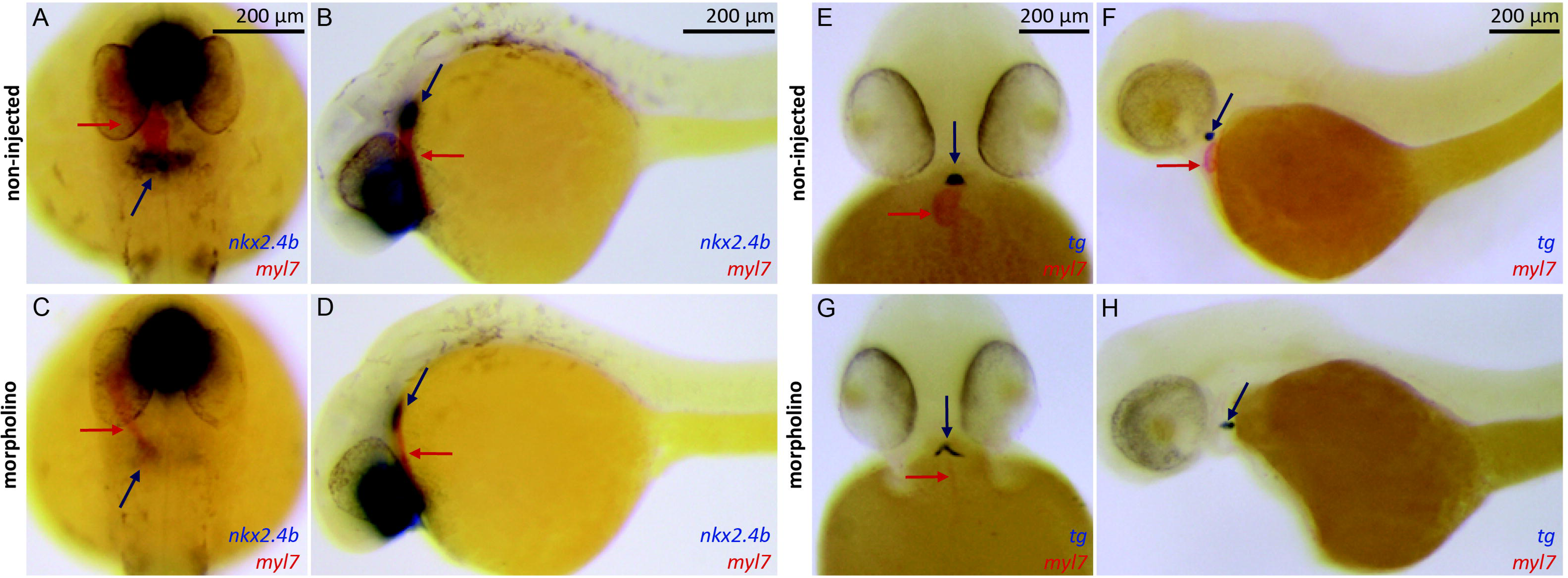
Impaired cardiac development in mef2c/d morpholino-injected embryos is associated with reduced thyroid marker expression. (**A-D**) Dual-color whole-mount *in situ* hybridization analysis of mRNA expression of markers for early thyroid (*nkx2.4b*) and cardiac (*myl7*) development in 28 hpf embryos injected with mef2c/d morpholino (MO) and non-injected control embryos (NI). MO injection caused variable loss of differentiated cardiomyocytes and resulted in smaller hearts (**C**) compared to controls. Concurrently, these embryos displayed reduced staining of *nkx2.4b* (**C**). (**E-H**) Dual-color whole-mount *in situ* hybridization analysis of mRNA expression of *myl7* and the thyroid differentiation marker *tg* in 55 hpf embryos injected with mef2c/d morpholino (MO) and non-injected control embryos (NI). MO-injected embryos displayed small, string-like midline hearts (**G**) and a modest reduction in *tg* staining (**G**). Blue arrows point to thyroidal *nkx2.4b* and *tg* expression and red arrows to *myl7* expression in the heart tube. Dorsal views (**A,C)** with anterior to the top, lateral views (**B,D,F,H**) with anterior to the left and ventral views (**E,G)** with anterior to the top are shown. Scale bar, 200 µm.

### Enhancing BMP signaling causes partial rescue of thyroid defects in BIO-treated embryos

Cardiac mesoderm expresses a number of diffusible growth factors, including several FGF and BMP ligands (17), (52) with potential thyroid specification-inducing capacity (17), (57), (58). In this respect, our recent small molecule screening showed that pharmacological inhibition of either FGF or BMP signaling during somitogenesis causes a failure of thyroid anlage formation in zebrafish (21). Thus, we reasoned that reduced availability of cardiac-borne signaling cues might explain the defective thyroid anlage formation observed in our experimental models (Wnt overaction, *mef2c/d*-morphants) displaying perturbed cardiac development in this study.

To address the question that diminished BMP signaling might contribute to thyroid phenotypes observed in embryos with after Wnt overactivation, we first examined endogenous *bmp4* expression in control embryos and embryos treated with BIO from 6 to 10 hpf. WISH analyses of 26 hpf embryos showed that the *bmp4* expression of cardiac tissue adjacent to the thyroid anlage region in normally developing embryos (**Supplementary figure 5A**) is completely ablated in BIO-treated embryos (**Supplementary figure 5B**). Similarly, Hinits *et al.* reported that Mef2c-deficient embryos lack *bmp4* expression in the cardiac region (52). In normally developing 42 hpf embryos, we also observed that cells contained in the forming thyroid bud display enhanced BMP signaling compared to adjacent foregut tissue (**Supplementary figure 5C-E**).

Next, we used a heat shock-inducible system to globally enhance BMP signaling during somitogenesis stages and assessed the effects of ectopically induced BMP signaling on thyroid development in control and BIO-treated embryos. For this purpose, we treated embryos from *Tg(hsp70l:bmp2b)* founders with either DMSO (0.1%) or BIO (5 µM) from 8 to 11 hpf and then exposed these embryos to heat-shock treatment at either early somitogenesis (11 hpf), mid-somitogenesis (15 hpf), late somitogenesis (20 hpf) or repeatedly during the course of somitogenesis (at 11, 15, 20 hpf). Subsequent WISH analyses of thyroid and cardiac markers at 28 hpf revealed different phenotypes depending on the pretreatment (DMSO, BIO) and the timing of heat shock-induced *bmp2b* overexpression.

Consistent with a proposed role of BMP for thyroid cell specification, global overactivation of BMP signaling in DMSO-treated embryos (controls) resulted in enhanced expression of *nkx2.4b* (**Figure 9A-I**). Not only was *nkx2.4b* staining strongly increased in these embryos, but the majority of affected embryos also showed a dramatic and irregular expansion of thyroid marker expression along the anterior-posterior axis. Concurrently, DMSO-treated embryos, when heat-shocked at early somitogenesis, failed to form a normal heart tube but showed a severely perturbed expression pattern of the cardiac marker *myl7* with bilateral expression domains extending irregularly along the anterior-posterior axis (**Figure 9C**). When we applied heat-shock on *Tg(hsp70l:bmp2b)* embryos at mid or late somitogenesis stages, we observed that the thyroid anlage and primitive heart tube formed at orthotopic positions (**Figure 9D-G**). We also observed a moderate enlargement of the *nkx2.4b* expression domain in these embryos. Embryos that were repeatedly heat-shocked during somitogenesis showed a phenotype similar to early somitogenesis heat shock (**Figure 9H,I**). Collectively, the timed heat-shock experiments in DMSO-treated embryos showed that the endoderm is competent at all somitogenesis stages to respond to BMP overactivation with an enhanced thyroid marker expression.

**Fig. 9.**
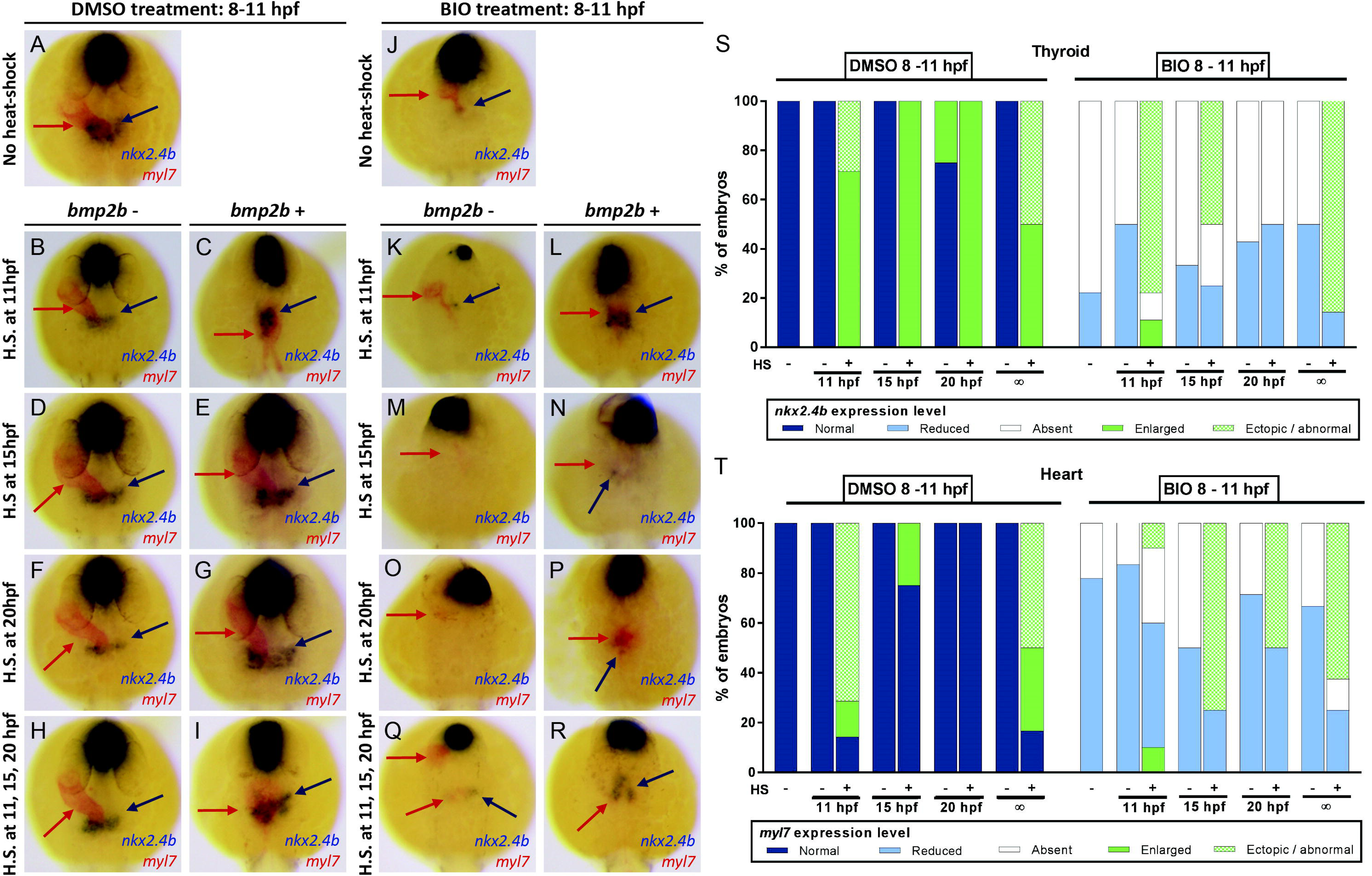
Conditional overactivation of BMP signaling in *Tg(hsp70l:bmp2b)* embryos partially rescues BIO-induced thyroid specification defects. (**A-R**) Dual-color whole-mount *in situ* hybridization analysis of thyroid (*nkx2.4b*) and cardiac (*myl7*) markers in 28 hpf embryos carrying the HS-inducible *bmp2b* transgene (*bmp2b*+) and in non-transgenic siblings (*bmp2b-*). Panels **A-I** show results for embryos that were treated with 0.1% DMSO (vehicle control) from 8-11 hpf and were subsequently exposed to timed heat shock (HS) treatment at early (11 hpf), mid (15 hpf) or late somitogenesis (20 hpf) and for embryos exposed to repeated HS at 11, 15 and 20 hpf. Carriers of the *bmp2b* transgene showed enhanced *nkx2.4b* expression in response to HS irrespective of the specific timing of HS treatment. Note that HS at 11 hpf (**B,C**) and repeated HS treatment (**H,I**) disrupted heart formation and caused ectopic stretches of *myl7* expression bilateral to the midline. Panels **J-R** show corresponding effects of HS treatment for embryos that were treated with 5 µM BIO from 8-11 hpf. HS of non-trangenic embryos did not affect the BIO-induced deficits in thyroid and cardiac development (compare **J** to **K,M,O,Q**). Carriers of the *bmp2b* transgene showed variable levels of *nkx2.4b* expression rescue, depending on the specific timing of HS treatment. Note that HS at 11 hpf (**K,L**) and the repeated HS treatment (**Q,R**) were most effective in restoring robust *nkx2.4b* expression. Partial recovery of *myl7* expression was detectable in response to *bmpb2b* overexpression, but heart tube formation was never restored. Dorsal views with anterior oriented to the top are shown. Blue arrows point to *nkx2.4b* expression in the thyroid anlage and red arrows to domains of *myl7* expression. Scale bar: 200 µm. (**S,T**) Quantification of the proportion of specimens displaying thyroid (**S**) and cardiac phenotypes (**T**) in response to different HS treatments as determined by *nkx2.4b* (**S**) and *myl7* (**T**) staining of 28 hpf embryos. Results are presented as the percentage of embryos displaying a particular phenotype. The total number of specimens analyzed for each treatment group is provided on the top of each column.

Heat-shock experiments with embryos pretreated with BIO (5 µM) from 8-11 hpf showed that global overactivation of BMP signaling could partially rescue the BIO-induced lack of thyroid marker expression at 28 hpf. WISH analyses and subsequent genotyping of stained specimens showed that heat-shock treatment of non-transgenic BIO-treated embryos did not improve *nkx2.4b* expression, irrespective of the timing of heat-shock induction during somitogenesis (**Figure 9J-R**). In stark contrast, BIO-treated embryos carrying the *hsp70l:bmp2b* transgene showed a strong *nkx2.4b* expression in the thyroid region when heat-shock induction of BMP signaling was performed at early somitogenesis (11 hpf) (**Figure 9K,L**) or when heat-shock was repeatedly applied at 11, 15 and 20 hpf (**Figure 9Q,R**). Just as observed in DMSO-treated embryos **(see** **Figure 9C**), overactivation of BMP signaling in BIO-treated embryos at early somitogenesis resulted in irregularly shaped and positioned *nkx2.4b* expression domains, often characterized by an elongated shape along the anterior-posterior axis (**Figure 9L**).

The capacity of BMP overactivation to rescue thyroid marker expression in BIO-treated embryos decreased markedly at later somitogenesis stages **(see** **Figure 9N**), and no rescue of *nkx2.4b* expression was detectable for heat-shock treatments carried out at 20 hpf **(see** **Figure 9P**). Thus, in contrast to DMSO-treated embryos, embryos treated with BIO lacked the competence to enhance thyroidal *nkx2.4b* expression in response to overactivation of BMP signaling at late somitogenesis stages. Accordingly, in this model, enhancing BMP signaling alone at mid- and late somitogenesis stages is not sufficient to rescue thyroid specification defects.

One characteristic feature of BIO-treated embryos displaying a rescue in *nkx2.4b* expression following early heat-shock treatment (11 hpf) was the presence of irregular ectopic patches of *myl7*-expressing cells near the *nkx2.4b*-expressing cells. Although we observed variable amounts of *myl7*-expressing cells across individual embryos from all heat-shock treatment groups **(see** **Figure 9N,P**), it was only in the group of BIO-treated embryos receiving heat-shock treatment at early somitogenesis (11 hpf) that we consistently observed a surplus of *myl7*-expressing cells in the vicinity of *nkx2.4b*-expressing cells. However, we note that we never observed a near full rescue of heart tube formation due to our heat shock treatments in BIO-treated embryos.

### Enhancing BMP signaling causes partial rescue of thyroid defects in Mef2c-deficient embryos

Given that enhanced BMP signaling could only achieve a rescue of thyroid specification defects in BIO-treated embryos under conditions that concurrently induced a small but consistent cardiac cell differentiation, we next applied a similar rescue approach to Mef2c-deficient. The Mef2c*-*deficient model provides a promising alternative rescue scenario as global BMP overactivation was deemed unlikely to overcome the block of cardiomyocyte differentiation resulting from the deficiency of mef2 protein function. In these experiments, we injected embryos from *Tg(hsp70l:bmp2b)* founders with *mef2c/d-*MO or maintained *Tg(hsp70l:bmp2b)* embryos as a non-injected control group. Injected and non-injected embryos were then heat-shock treated at early (10 hpf), mid (15 hpf), or late somitogenesis (20 hpf) and thyroid and cardiac markers were analyzed in the different treatment groups at 28 hpf (**Figure 10**). In the non-injected control embryos, we observed changes in thyroid and cardiac development (**Figure 10A-G**) that were very similar to the effects seen in DMSO-treated control embryos in the previous experimental series (**Figure 9A-G**). Depending on the timing of heat-shock treatment, we observed ectopic expansions of *nkx2.4b* expression along the anterior-posterior axis (heat-shock at 10 hpf) or moderately enhanced *nkx2.4b* expression when heat-shock was applied at mid (15 hpf) or late somitogenesis stages (20 hpf). In addition to the irregular shape and position of the thyroidal *nkx2.4b* expression domain, we noted again abnormal *myl7* expression domains, irregularly extending along the anterior-posterior axis (**Figure 10C**).

**Fig. 10.**
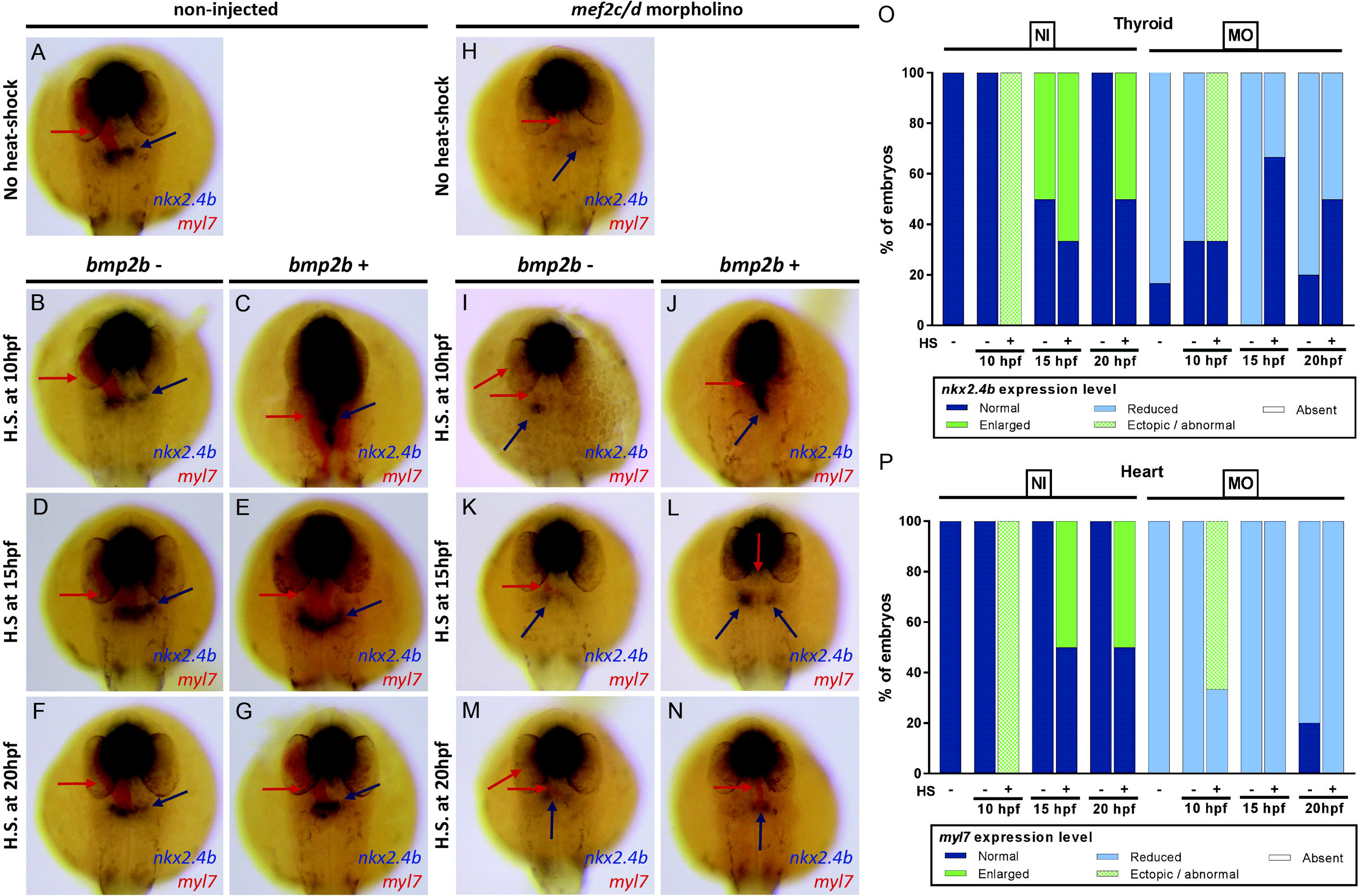
Conditional overactivation of BMP signaling in *Tg(hsp70l:bmp2b)* embryos partially rescues mef2c/d morpholino-induced thyroid specification defects. (**A-N**) Dual-color whole-mount *in situ* hybridization analysis of thyroid (*nkx2.4b*) and cardiac (*myl7*) markers in 28 hpf embryos carrying the HS-inducible *bmp2b* transgene (bmp2b+) and in non-transgenic siblings (bmp2b-). Panels **A-G** show results for control embryos exposed to timed heat shock (HS) treatment at early (10 hpf), mid (15 hpf) or late somitogenesis (20 hpf). Carriers of the *bmp2b* transgene showed enhanced *nkx2.4b* expression in response to HS irrespective of the specific timing of HS treatment. Note that HS at 10 hpf (**B,C**) disrupted heart formation and caused ectopic stretches of *myl7* expression bilateral to the midline. Panels **H-N** show corresponding effects of HS treatment for embryos that were injected with mef2c/d morpholino (MO) at the one-cell stage. HS of non-trangenic embryos did not affect the MO-induced deficits in thyroid and cardiac development (compare **H** to **I,K,M**). Carriers of the *bmp2b* transgene showed variable levels of *nkx2.4b* expression rescue, depending on the specific timing of HS treatment. Note that HS at 10 hpf (**I,J**) was most effective in restoring robust *nkx2.4b* expression. No significant recovery of *myl7* expression was detectable in response to *bmpb2b* overexpression. Dorsal views with anterior oriented to the top are shown. Blue arrows point to *nkx2.4b* expression in the thyroid anlage and red arrows to domains of *myl7* expression. Scale bar: 200 µm. (**O,P**) Quantification of the proportion of specimens displaying thyroid (**O**) and cardiac phenotypes (**P**) in response to different HS treatments as determined by *nkx2.4b* (**O**) and *myl7* (**P**) staining of 28 hpf embryos. Results are presented as the percentage of embryos displaying a particular phenotype. The total number of specimens analyzed for each treatment group is provided on the top of each column.

WISH analyses of the thyroid and cardiac marker expression of Mef2c*-*deficient embryos at 28 hpf showed a marked reduction in the thyroid *nkx2.4b* staining (**Figure 10H**). The severity of the thyroid specification defects correlated with the severity of a concurrent cardiac developmental phenotype, as judged from the expression of *myl7*. As expected, irrespective of their timing, heat-shocks of non-transgenic Mef2c*-*deficient embryos failed to rescue the thyroid or cardiac phenotype (**Figure 10I,K,M**). In Mef2c*-*deficient embryos carrying the *hsp70l:bmp2b* transgene, heat-shock induction of BMP signaling at early somitogenesis (10 hpf) resulted in a robust thyroid *nkx2.4b* expression, even though the thyroid’s shape and position were abnormal (**Figure 10J**). This phenotype largely resembled the expansion along the anterior-posterior axis seen in control embryos heat-shocked at 10 hpf (**Figure 10C**). Similar to the series of heat-shock experiments performed on BIO-treated embryos, a partial rescue of thyroid specification defects was generally limited to heat-shock induction of BMP signaling at early somitogenesis stages. In Mef2c-deficient embryos carrying the *hsp70l:bmp2b* transgene, heat-shock induction of BMP signaling at mid-(15 hpf) or late somitogenesis (20 hpf) only mildly increased levels of *nkx2.4b* expression compared to untreated *mef2c/d* morphants (**Figure 10L,N**). Compared to the thyroid rescue observed in BIO-treated embryos, a notable difference in the Mef2c-deficient model was that heat-shock induction of BMP signaling did not rescue the cardiac differentiation phenotype. Thus, our observations in the cardiac deficient *mef2c/d* morphant model indicate that ectopically induced BMP signaling is sufficient to rescue the thyroid specification defect even in the absence of cardiomyocyte differentiation.

## Discussion

In this study, we described severe defects in early zebrafish thyroid development as a result of globally modulated canonical Wnt signaling during gastrula and early somitogenesis stages. While thyroid dysgenesis in response to drug-induced Wnt signaling modulation was first detected in the course of our recent small molecule screening study with zebrafish embryos (21), the current study provides corroborative evidence that the thyroid specification defects observed following treatment with BIO and AZA are due to the up-regulation of canonical Wnt signaling. We used a well-characterized zebrafish biosensor line (35) to confirm that BIO and AZA treatments are effectively inducing canonical Wnt signaling *in vivo* during zebrafish development. Our results are, therefore, in line with numerous studies in zebrafish, *Xenopus,* and mammalian cell systems utilizing these two compounds to globally induce Wnt signaling (35), (59), (60). Importantly, the thyroid specification defects observed in drugged embryos were faithfully reproduced in a genetic zebrafish, thereby corroborating critical evidence that the thyroid defects are indeed caused by enhanced Wnt signaling and are not due to possible non-specific side effects of drug treatments.

Thyroid phenotyping of drugged embryos at the thyroid anlage stage (28 hpf), using *nkx2.4b* expression as a proxy for thyroid lineage commitment, revealed concentration-dependent losses of *nkx2.4b*-expressing cells in the prospective thyroid region. In zebrafish embryos, thyroidal expression of *nkx2.4b* typically becomes detectable at around 23/24 hpf, and a global assessment of developmental timing in drugged embryos indicated that reduced detection of *nkx2.4b* expression was not solely due to delayed development. Remarkably, gastrula treatment with a high concentration (5 µM) of BIO or AZA caused a complete absence of detectable *nkx2.4b* expression in 28 hpf embryos. Although the latter phenotype was highly penetrant at 28 hpf (affecting almost 100% of drugged embryos), analyses of treated embryos at later stages (55 hpf) showed that almost all treated embryos developed at least a very tiny thyroid primordium comprised of very few cells expressing the early functional thyroid marker *tg*. It is currently unknown if these few remaining *tg*-expressing thyroid cells at 55 hpf are derived from thyroid precursors expressing *nkx2.4b* at such low levels that are undetectable by our WISH at 28 hpf or if some precursor cells are specified after 28 hpf to give rise to *tg*-expressing thyroid cells. Despite the presence of very few differentiated thyroid cells at later stages, we can conclude that increased canonical Wnt signaling during early zebrafish development impairs the specification of thyroid precursor cells.

By progressively shifting the timing of short-term drug treatments, we observed that the primary events leading to this thyroid dysgenesis phenotype occur during gastrula and early somitogenesis stages. Therefore, the question arises whether increased Wnt signaling directly affects development of endodermal progenitors, which later give rise to foregut endoderm and the thyroid cell lineage, or whether indirectly through another cell type (e.g. pre-cardiac mesoderm) that is required later in development as a local source of signals to initiate thyroid specification in the foregut endoderm zebrafish (18), (27).

During early vertebrate development, Wnt signaling plays a key role in the anteroposterior patterning, mainly by acting as a posteriorizing factor (61). This anterior-posterior patterning role of Wnt signaling has been extensively studied during neural development (55). In our studies, a robust posteriorizing activity of Wnt on neural development was evident from the severe forebrain and eye development defects. Since the thyroid develops from the anterior foregut endoderm, the posteriorizing activity of Wnt on the endoderm could provide a plausible hypothesis to explain the observed thyroid dysgenesis. Surprisingly few studies have addressed the precise role of Wnt on endoderm patterning. Studies in *Xenopus* embryos indicated that canonical Wnt signaling during gastrulation might pattern the endoderm in a way much similar to what is known for the nervous system (27). Notably, forced Wnt8 expression in cells fated to become endoderm was found to block anterior endoderm development in *Xenopus* embryos, resulting in specification failure of foregut organ primordia. In contrast to the aforementioned *Xenopus* studies, morphological and molecular analyses of thyroid-lacking zebrafish embryos revealed the formation of a morphologically fairly normal anterior endoderm with a foregut identity (*foxa2* expression) and confirmed timely expression of various other foregut organ markers (*hhex*, *pdx1*, *prox1a*). Moreover, data presented by McLin *et al*. (27) indicate that thyroid specification was preserved in *Xenopus* embryos if enhanced β-catenin signaling was cell-autonomously restricted to the endoderm lineage. When comparing these study results with our observations, we conclude that posteriorization effects of BIO-induced Wnt activity were by far not as dramatic as reported in the *Xenopus* studies and that posteriorization of anterior endoderm patterning was likely not the primary cause for the severe thyroid specification defects in zebrafish embryos.

Another striking effect of drug-induced Wnt activation was a severe impairment of cardiac differentiation. Consistent with previous studies, we confirmed that enhanced Wnt signaling during zebrafish gastrulation in zebrafish limits cardiac differentiation within pre-cardiac mesoderm (25), (26). Intriguingly, we observed a close correlation between the disruption of cardiac progenitor differentiation and the thyroid abnormalities. This was true for experiments involving increasing concentrations of BIO and AZA as well as for experiments with different treatment periods. In this respect, it is noteworthy that treatment periods, which caused little if any global posteriorization effects, still showed reduced cardiac differentiation concurrent with thyroid specification defects. Thus, the apparent relationship between the diminished formation of cardiac tissue and the corresponding defects in thyroid specification led us to formulate the hypothesis that the primary effect of enhanced Wnt signaling might be the blockage of cardiac differentiation while the thyroid defects might be a secondary event resulting from a reduction of cardiac-borne signaling cues. This hypothesis is supported by results from several previous studies in different vertebrate models, which provided evidence for a critical role of pre-cardiac and cardiac mesoderm in the induction of thyroid precursor cells signaling (17), (18).

Our own studies provided further supporting lines of evidence for this hypothesis by showing that a similar correlation between impaired cardiogenesis and defective thyroid specification exists in Mef2c-deficient embryos (52), a model in which heart formation is blocked independent of perturbations in Wnt signaling. Since the mesoderm, and not the foregut endoderm, expresses *mef2ca* and *mef2cb*, this model provided more direct evidence for the contention that thyroid specification relies on proper differentiation of cardiac tissue. A role of mesoderm as a source of signals coming to induce specification of thyroid progenitors within the foregut endoderm is also consistent with what has been already described concerning the specification of other endoderm-derived organs (62), (63), (64).

While these observations strongly suggest an important role of cardiac mesoderm as a signal source involved in the early steps of thyroid development, it raises the question of the nature of the signaling cues involved in this tissue-tissue interaction. Based on results from our small molecule screening (21) and previous studies in several other model systems, BMP signaling was deemed a primary candidate of particular importance for thyroid specification (65). *In vitro* experiments with murine stem cell models identified key conserved roles for BMP signaling in regulating thyroid lineage specification from foregut endoderm (65). On the other hand, Wnt was not required for thyroid specification from endoderm *in vitro*, like it had been previously suggested (66). Expression profiles during embryogenesis also support a potential role of BMP signaling as a key factor in regulating early steps of thyroid development. In mouse, BMP4 is the Bmp ligand expressed at high levels by the cardiac mesoderm, in particular within the secondary heart field, the structure that gives rise among others to the outflow tract, which the developing thyroid lies nearby (67), (68).

Danesh *et al.* also demonstrated in mouse embryos that BMPR1a is highly expressed in regions of the pharyngeal endoderm, where endodermal progenitors will differentiate into thyroid cells (68). Thus, Bmp4 could be one critical endogenous Bmp ligand acting from mesoderm to endoderm to initiate thyroid development through interaction with its receptor Bmpr1a. For zebrafish, we further confirmed a high *bmp4* expression in cardiac mesoderm around the time of thyroid specification (Supplementary Figure 5) and ectopic BMP signaling during somitogenesis caused an expansion of the thyroid anlage (**Figures 9 & 10**).

Against this background, it is noteworthy that timed overactivation of BMP signaling during zebrafish somitogenesis stages could partially restore thyroid specification in both models of impaired cardiac development. Interestingly, in our model of cardiac dysgenesis due to enhanced Wnt signaling, the restoration of thyroidal *nkx2.4b* expression achieved by subsequent BMP overactivation appeared strictly dependent on a concurrent rescue of cardiac cell differentiation. At this stage, it is therefore not possible to conclude if thyroid specification was directly restored by the ectopic BMP activity in this model. Several lines of evidence indicate that thyroid specification requires the combined action of BMP and FGF signaling (17), (18), (31), (65), (57). One possible interpretation might be that BMP signaling alone might not be sufficient to induce thyroid specification in embryos with almost no cardiac mesoderm formation. In contrast, restoration to some extent of cardiac differentiation by transient BMP overactivation could provide a source of endogenous BMP, FGF, and possibly other signaling cues to induce thyroid specification.

Rescue of thyroid specification was also evident in Mef2c-deficient embryos after BMP overactivation. However, in this model, the correlation between thyroid rescue and the presence of cardiac mesoderm was less explicit. This correlation reduction was due to the incomplete penetrance of the cardiac phenotype in morphant embryos. However, we observed many Mef2c-deficient embryos presenting a partial thyroid rescue while still lacking cardiomyocytes. These observations indicate that ectopic BMP activation might indeed restore thyroid specification even in the absence of detectable cardiomyocytes. To understand the differences between the thyroid rescue induced by ectopic BMP activity in the two cardiac dysgenesis models, the different mechanisms leading to deficient cardiomyocyte development need to be taken into account. Enhanced Wnt signaling at early stages leads to a block of early cardiac mesoderm differentiation with a pronounced deficit in cardiac progenitor differentiation (see Figure 5). In contrast, studies by Hinits *et al* (52) showed that early cardiac development is uncompromised in Mef2c-deficient embryos and that the last steps involved in cardiomyocyte differentiation are blocked in this model, including the up-regulation of *bmp4* expression in the cardiac field. Thus, pre-cardiac mesoderm present in Mef2c-deficient embryos might serve as a source of FGF and other signaling factors needed to act in concert with BMP to induce thyroid specification. Given that Mef2c-deficient embryos lack *bmp4* expression in the cardiac mesoderm and that ectopic overexpression of *mef2cb* mRNA can induce *bmp4* in the cardiac field (52), it would be interesting to investigate further this model, respect to the expression of other potential thyroid-inducing pathways (e.g., FGF) and their role in acting together with BMP signaling in the process of thyroid specification. Also, results obtained in Mef2c-deficient embryos indicate that the thyroid of Bmp4-deficient embryo models might bring further insights into the role of BMP signaling in thyroid specification.

In summary, our studies confirmed and validated our previous results from a small molecule screen that enhanced Wnt signaling during gastrulation stages results in severe inhibition of thyroid specification. Moreover, we provide corroborating evidence for the contention that this thyroid dysgenesis results from a blockage of cardiac development and, consequently, the lack of instructive cardiac-derived signaling cues for the specification of thyroid precursor cells. This hypothesis is supported by a similar perturbation of thyroid specification in Mef2c-deficient embryos displaying a lack of cardiomyocyte differentiation. Finally, we demonstrate that ectopic activation of BMP signaling can partially rescue these thyroid phenotypes identifying BMP signaling as one critical component of this tissue-tissue interaction.

## Acknowledgments

We thank J.-M. Vanderwinden from the Light Microscopy Facility for technical assistance. This work was supported by grants from the Belgian National Fund for Scientific Research (FNRS) (FRSM 3-4598-12; CDR-J.0145.16), the Action de Recherche Concertée (ARC) de la Communauté Française de Belgique (ARC AUWB-2012-12/17-ULB3), the Fonds d’Encouragement à la Recherche de l’Université Libre de Bruxelles (FER-ULB), the Fund Yvonne Smits (King Baudouin Fundation) and the Berlin Institute of Health (BIH, CRG-TP2).

This work was supported by grant the Belgian National Fund for Scientific Research (FNRS): I.V. is FNRS Research Fellow, B.H. and P.G. are Fund for Research in the Industry and the Agriculture (FRIA) research fellow, R.O. was FNRS Postdoctoral fellow and S.C. is FNRS Senior Research Associate. We greatly appreciate the help of J. Bakkers, E. Moro, S. Abdelilah-Seyfried and G. Weidinger for providing us with transgenic lines used in this study. J. Bakkers for Tg(kdrl:EGFP) and Tg(myl7:EGFP) lines, E. Moro for *Tg(7xTCF-Xla.Siam:GFP)*, S. Abdelilah-Seyfried For *Tg(hsp70l:bmp2b) line, G. Weidinger For Tg(hsp70l:wnt8a-EGFP*)

## Corresponding author

Please address all correspondences and requests for reprints to: Sabine Costagliola, Institut de Recherche Interdisciplinaire en Biologie Humaine et Moléculaire, Université Libre de Bruxelles, 808 Route de Lennik, 1070 Brussels, Belgium.

Robert Opitz, Institute of Experimental Pediatric Endocrinology, Charité Universitätsmedizin Berlin, Augustenburger Platz 1, 13353, Berlin, Germany.

