## Supplementary Information for "Enhanced canonical Wnt signaling during early zebrafish development perturbs the interaction of cardiac mesoderm and pharyngeal endoderm and causes thyroid specification defects"

**This file contains:**

**Supplementary Table 1.** Gene-specific primers used for PCR-based approach of RNA probe preparation

**Supplementary Table 2.** Antibodies used in whole-mount *in situ* hybridization (WISH) and immunofluorescence (WIF) staining.

**Supplementary Figure 1.** Ectopic localizations of thyroid tissue following canonical Wnt inhibition during gastrula, in 28 & 55 hpf embryos.

**Supplementary Figure 2**. Wnt activity revealed by whole-mount in situ hybridization of *egpf* in Wnt-reporter line *tg(7xTCF-Xla.Siam:GFP)ia4* during different Wnt chemical modulations between 6 and 10 hpf.

**Supplementary Figure 3**. *Sox17* positive endodermal cells in BIO-treated embryos during gastrula in comparison to controls, in 30 hpf embryos.

**Supplementary Figure 4.** Dose-dependent impaired cardiac development following chemical canonical Wnt activation during gastrula (6-10 hpf), in 48 hpf embryos.

**Supplementary Figure 5**. *Bmp4* expression in cardiomyocytes around the timing of thyroid specification and Bmp signaling activation in early thyroid development.

**Supplemental Table 1**

Gene-specific primers used for PCR-based approach of RNA probe preparation

| Gene symbol | GenBank Accession | Gene-specific forward primer 5’ → 3’ | Gene-specific reverse primer 5’ → 3’ | RNA probe size (bp) |
|---|---|---|---|---|
| ***nkx2.4b*** | NM_131589 | TTTCTCAGTGAGCGATATTTGAG | CGTATACAATTCAGTCAAAGAGG | 1058 |
| ***tg*** | NM_001329865 | TAATTGTAGTGACGGCCAGTTTTA | TTTTCTCCTTGTAGCTGAAGAGGT | 972 |
| ***nkx2.5*** | NM_131421 | ATGTTCTCTAGCCAAATGACTTCC | TATTTAGATCCCCAACACCAAAGT | 877 |
| ***gfp*** | L29345 | GACGTAAACGGCCACAAGTTCAGC | CTTCTCGTTGGGGTCTTTGCTCAG | 582 |
| ***mef2cb*** | NM_001130962 | AAAAAGATTCAGATCACACGGATT | GAGGGCCATTGCTTATCATGTG | 1434 |
| ***gata4*** | NM_131236 | TAATCGCTCACCTGTGGATATGTA | GCTGTTCCACACTTCACTCTTG | 1021 |
| ***gata5*** | NM_131235 | AGTTTTTCGACTGTCGCTTATTTT | CATGTCTATACTGGACCTCCTTCC | 814 |
| ***foxa2*** | NM_130949 | ATCCGAGTCGAAAAACATCAGAG | CATGTGATTCAAGAAGTCCAATC | 677 |
| ***foxa3*** | NM_131299 | AGCTCCAAATCTTTTACACAGCTC | TTTGCGTACACCTTTCTGTTACAT | 760 |
| ***hhex*** | NM_130934 | TATTGAAGACATCTTGGGAGAAC | TCCCCAAAATATCCTTTATTGAA | 756 |
| ***pdx1*** | NM_131443 | CCTAACCACCTGTACAAGGACTCT | CATAACCTTCAAACAATGTCCAAA | 874 |
| ***prox1a*** | NM_131405 | CTCAGCTTCTGAAAAACAATTTGA | CACCTGCATACTACCAAAAGTGTC | 1000 |
| ***bmp4*** | NM_131342 | ACTATCCTGAAAGATCCACCAGTC | TCGATCAGACAAGAAATGACAAA | 990 |

Note: A sequence containing the T3 polymerase promoter was added to the 5’end of each reverse primer sequence in order to obtain a PCR product that could be used to generate an antisense probe during in vitro transcription. Therefore, the final design of the reverse primers used in this study was as follows: 5’-GGATCCAATTAACCCTCACTAAAGGGAA (N)24-3’ with N representing the 24 nucleotides of the gene-specific reverse primer. The bases of the T3 polymerase promoter become double-stranded promoter sequence during the PCR reaction.

**Supplemental Table 2**

Antibodies used in whole-mount *in situ* hybridization (WISH) and immunofluorescence (WIF) staining.

| Antibody | Dilution | Host | Supplier (Cat.) | WIF/WISH |
|---|---|---|---|---|
| anti-DIG antibody conjugated to alkaline phosphatase | 1:6000 | sheep | Roche (11093274001) | WISH |
| anti-fluorescein antibody conjugated to alkaline phosphatase | 1:2000 | sheep | Roche (1426338) | WISH |
| anti-DNP antibody conjugated to alkaline phosphatase | 1:600 | rabbit | Vector Laboratories (MB-3100) | WISH |
| anti-DIG antibody conjugated to horseradish peroxidase | 1:500 | sheep | Roche (11207733910) | WISH |
| anti-GFP polyclonal antibody | 1:1000 | chicken | Abcam (ab13970) | WIF |
| anti-DsRed polyclonal antibody | 1:250 | rabbit | Clontech (632496) | WIF |
| Alexa Fluor488-conjugated anti-chicken IgG antibody | 1:250 | goat | Invitrogen (A11039) | WIF |
| Alexa Fluor488-conjugated anti-rabbit IgG antibody | 1:250 | donkey | Jackson ImmunoResearch (711-545-152) | WIF |
| Cy3-conjugated anti-rabbit IgG antibody | 1:250 | donkey | Jackson ImmunoResearch (711-165-152) | WIF |
| Alexa Fluor647-conjugated anti-mouse IgG antibody | 1:250 | donkey | Jackson ImmunoResearch (715-605-150) | WIF |

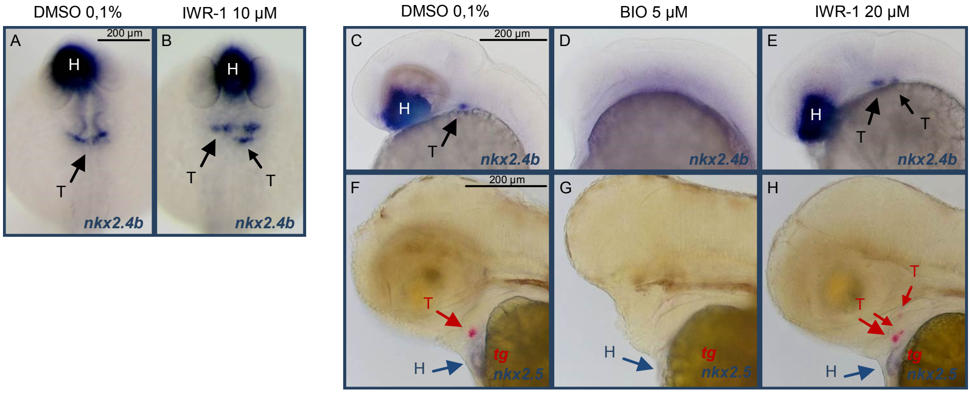


**SUPPLEMENTARY FIGURE 1:**

**Ectopic localizations of thyroid tissue following canonical Wnt inhibition during gastrula, in 28 & 55 hpf embryos.**

Panels (A) and (B) show dorsal view, anterior oriented to the top, and panels (C) to (H) show lateral view, anterior oriented to the left, of embryos.

(A) and (B): Expression of the early thyroid marker *nkx2.4b* in 28 hpf embryos following treatment with 0.1% DMSO (vehicle control) and Wnt-inhibiting drugs, IWR-1, between 6 and 10 hpf. In control embryos (A), *nkx2.4b* is expressed in the thyroid anlage (T) and the ventral forebrain (hypothalamus, H). In response to IWR-1 treatment (B), *nkx2.4b* mRNA expression is broader, with relative ectopic localization (small arrow). Arrows point to *nkx2.4b* expression in the thyroid anlage.

(C) to (E): Expression of the early thyroid marker *nkx2.4b* in 28 hpf embryos following treatment with 0.1% DMSO (vehicle control), BIO Wnt-inducing drug and IWR-1 Wnt-inhibiting drug between 6 and 10 hpf. In control embryos (C), *nkx2.4b* is expressed in the thyroid anlage (T) and the ventral forebrain (hypothalamus, H). In response to BIO (D), *nkx2.4b* mRNA expression is lost in both domains. In response to IWR-1 treatment (E), *nkx2.4b* mRNA expression is broader, with relative ectopic localization (small arrow). Arrows point to *nkx2.4b* expression in the thyroid anlage.

(F) to (H): Expression of the thyroid (T, in red) marker *tg* and the heart (H, in blue) marker *nkx2.5* in 55 hpf embryos following treatment with 0.1% DMSO (vehicle control), BIO Wnt-inducing drug and IWR-1 Wnt-inhibiting drug between 6 and 10 hpf. In control embryos (F), *tg* is expressed in the thyroid anlage (T). In response to BIO (G), *tg* mRNA expression is lost and cardiac *nkx2.5* expression is decreased. In response to IWR-1 treatment (H), *tg* expression is broader, with ectopic localizations (smalls arrows). Red arrows point to *tg* expression in the thyroid anlage, blue arrows point to *nkx2.5* expression in the heart.

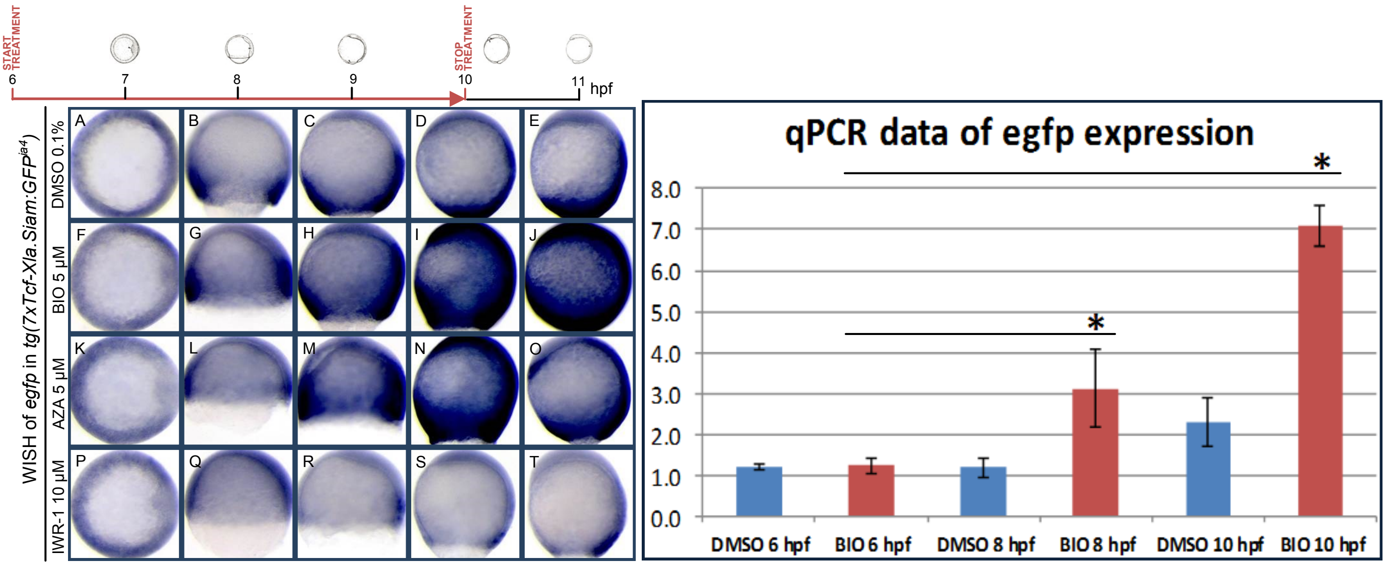


**SUPPLEMENTARY FIGURE 2:**

**Wnt activity revealed by whole-mount in situ hybridization of *egpf* in Wnt-reporter line *tg(7xTCF-Xla.Siam:GFP)*$^{ia4}$ during different Wnt chemical modulations between 6 and 10 hpf.**

Panels (A), (F), (K) and (P) show dorsal, anterior oriented to the left, and other panel show lateral views, anterior oriented to the top, of embryos.

(A) to (E): Intensity of *egfp* riboprobe staining one hour (A), two hours (B), three hours (C), four hours (D) in a control situation (DMSO incubation of embryos from 6 to 10 hpf) and one hour after removing DMSO from the medium (E).

(F) to (J): Intensity of *egfp* riboprobe staining one hour (F), two hours (G), three hours (H), four hours (I) after starting incubation with the chemical Wnt activator BIO with a concentration of 5 µM and one hour after stopping it (J), showing increase of *egfp* expression, corresponding to enhanced canonical Wnt activity.

(K) to (O): Intensity of *egfp* riboprobe staining one hour (K), two hours (L), three hours (M), four hours (N) after starting incubation with the chemical Wnt activator AZA with a concentration of 5 µM and one hour after stopping it (O), showing increase of *egfp* expression, corresponding to enhanced canonical Wnt activity.

(P) to (T): Intensity of *egfp* riboprobe staining one hour (P), two hours (Q), three hours (R), four hours (S) after starting incubation with the chemical Wnt inhibitor IWR-1 with a concentration of 10 µM and one hour after stopping it (T), showing decrease of *egfp* expression, corresponding to reduced canonical Wnt activity.

(U): qPCR data of *egfp* expression in controls (DMSO treatment) in comparison to those treated with BIO at 6 hpf, 8 hpf (2 hours after starting incubation with the chemical) and 10 hpf (4 hours after starting incubation with the chemical). Asterisks denote a statistically significant increase of *egfp* expression after 2 hours of BIO treatment and 4 hours of BIO treatment (compared to *egfp* expression in 6 hpf embryos).

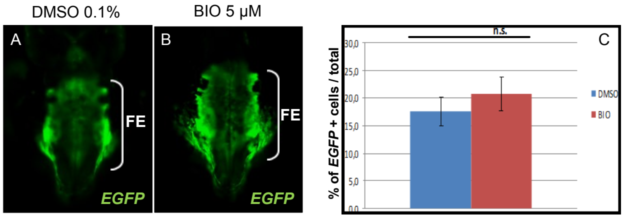


**SUPPLEMENTARY FIGURE 3:**

***Sox17* positive endodermal cells in BIO-treated embryos during gastrula in comparison to controls, in 30 hpf embryos.**

Panels show dorsal view of embryos, anterior oriented to the top.

(A) and (B): Whole-mount immunofluorescence of *tg(sox17:EGFP)* embryos with anti-*EGFP* antibodies at 30 hpf showing comparable endodermal pattern in a control situation / DMSO 0,1% treatment (A) and in embryos treated with 5 µM of BIO during gastrula (B). Brackets show *EGFP* expression in the endodermal region. FE = foregut endoderm.

(C): FACS counting of *EGFP* cells at 30 hpf shows no statistically significant difference of percentage of *EGFP* positive cells in control embryos in comparison to BIO-treated embryos during gastrula.

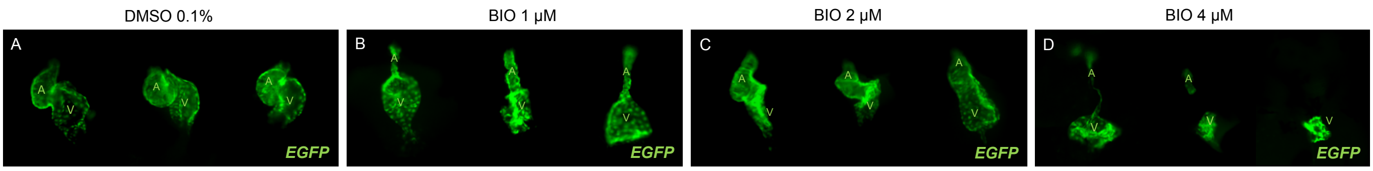


**SUPPLEMENTARY FIGURE 4:**

**Dose-dependent impaired cardiac development following chemical canonical Wnt activation during gastrula (6-10 hpf), in 48 hpf embryos.**

Panels show frontal view of embryos, anterior oriented to the top.

(A) to (D): Whole-mount immunofluorescence of *tg(myl7:EGFP)* embryos with anti-*EGFP* antibodies at 48 hpf showing normal heart tube in a control situation / DMSO 0,1% treatment in (A), and progressive reduction of cardiomyocytal *myl7* expression after gastrula BIO treatment with 1 μM (B), 2 μM (C) and 4 μM (D). Each panel shows heart of 3 different embryos. Images were taken by epifluorescence microscope. A = atrium. V = ventricle.

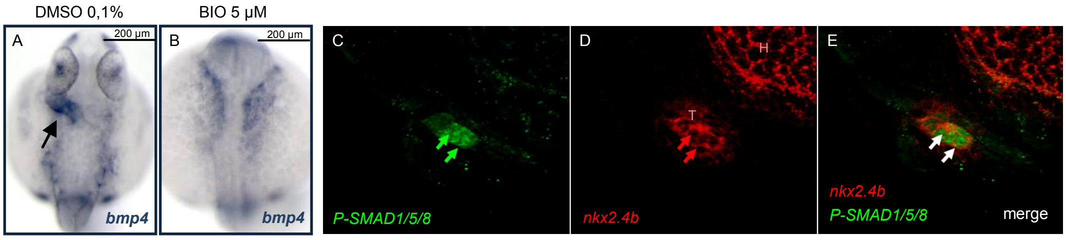


**SUPPLEMENTARY FIGURE 5:**

***Bmp4* expression in cardiomyocytes around the timing of thyroid specification and Bmp signaling activation in early thyroid development.**

Panels (A) and (B) show dorsal view of embryos, anterior oriented to the top. Panels (C) to (E) show lateral view of embryos, anterior oriented to the right, zoomed on the thyroid.

(A) and (B): Whole-mount in situ hybridization (WIF) of WT embryos at 26 hpf showing high *bmp4* expression in the cardiac tube in a control situation / DMSO 0,1% treatment in (A) and absence of such expression after gastrula (6-10 hpf) BIO treatment with 5 µM (B). Arrow points to *bmp4* expression in the cardiac tube.

(C) to (E): Whole-mount immunofluorescence (WISH) of WT embryos with anti-*P-SMAD1/5/8* antibodies at 42 hpf, just after the detachment of the thyroid from the pharyngeal floor, showing high Bmp activity in nuclei of thyroid cells in a control situation / DMSO 0,1% treatment (C). In panel D, the cytoplasm of thyroid cells is stained with *nkx2.4b* fluorescent WISH and in (E), both WIF and fluorescent WISH are merged. Images were taken by epifluorescence microscope. Green arrows point to *P-SMAD1/5/8* expression in nuclei of thyroid cells. Red arrows point to *nkx2.4b* expression in cytoplasm of thyroid cells. White arrows point to merged *P-SMAD1/5/8* and *nkx2.4b* expressions. T = thyroid primordium. H = hypothalamus.
